# The neural basis of flavour-nutrient learning

**DOI:** 10.1101/2025.02.09.637334

**Authors:** James C. R. Grove, Anna M. Hakimi, Queenie Li, Jingkun Zhang, Heiko Backes, Bojana Kuzmanovic, Jaewon Choi, Violeta Ubadiah, Longhui Qiu, Zhengya Liu, Dana M. Small, Marc Tittgemeyer, Zachary A. Knight

## Abstract

Animals learn about the external world, in part, via interoceptive signals^1,2^. For example, the nutrient content of food is first estimated in the mouth, in the form of flavour, and then measured again via slower signals from the gut. How these signals from the mouth and gut are integrated to drive learning is unknown. Here we identify a dopamine pathway that enables learning about the nutrient content of food. We show that dopamine neurons in the ventral tegmental area (VTA_DA_) are necessary for associating nutrients with flavours, and that post-ingestive nutrients trigger DA release selectively in a small region of the anterior basolateral amygdala (BLA) but not canonical DA targets in striatum. This nutrient-triggered DA release occurs preferentially in the left BLA in mice, and this same structure is preferentially activated by post-ingestive nutrients in humans. We identify the gut sensors that are responsible for nutrient-triggered DA release; show that they activate BLA-projecting DA neurons defined by expression of cholecystokinin (CCK); and demonstrate that stimulation of DA axon terminals in the anterior BLA drives flavour-nutrient learning but not other aspects of feeding behavior. Two-photon imaging of neurons in the left anterior BLA reveals that they integrate gustatory and post-ingestive cues, and silencing these neurons prevents flavour-nutrient learning. These findings establish a neural basis for how animals learn about the nutrient content of their food.

## Introduction

We learn about food during the process of eating^1,2^. For example, animals come to prefer calorie-rich foods after experiencing their post-ingestive effects^1,2^. This learning process is enabled by sensation along the alimentary canal that begins at the tip of the tongue and extends into the intestine. A fundamental challenge is to explain how these diverse interoceptive signals are integrated in the brain to create a coherent picture of what has been consumed.

Prior work has shown that there is a privileged connection between sensory signals that arise from the mouth and gut. For example, animals can learn to lick a flavoured solution, but not press a lever, in order to receive an infusion of nutrients into the stomach^3,4^. This suggests that there should be hardwired pathways in the brain for linking orosensory and gut signals.

VTA_DA_ neurons are important for learning relationships between cues and rewards^4,5^, and, while this has traditionally been studied in the context of exterosensory cues^5–7^, pharmacological studies suggest DA is also involved in associating post-ingestive nutrients and flavours^8–11^. For example, dopamine antagonists delivered to cortex^9^, striatum^10^, and amygdala^11^ can all partially impair this learning process. However, the specific contribution of these structures is unclear. We therefore set out to identify the cell types, pathways, and mechanisms involved.

### DA neurons are required for learning about nutrients but not toxins

We first tested whether VTA_DA_ neurons are required for learning associations between nutrients and flavours. We targeted the inhibitory opsin stGtACR to these cells for neural silencing, and, in a separate surgery, equipped mice with intragastric (IG) catheters for nutrient infusion. We then used a closed-loop paradigm for post-ingestive learning^3,12^ in which mice are given access to two differentially flavoured solutions, one of which is paired with lick-triggered IG glucose infusion and the other of which is paired with isovolumic infusions of water (see Methods, Fig. 1a). After six days of training, the preference of mice for these two flavours is measured in a two-bottle test. Consistent with previous findings^12^, we found that control mice developed a robust preference for the solution paired with sugar infusion (35±5% before vs. 81±7% after training, p=7.8e-5). In contrast, optogenetic silencing of VTA_DA_ neurons throughout training completely blocked the acquisition of a flavour preference (48±7% before vs. 53±7% after training, Fig. 1b; Extended Fig. 1). Importantly, silencing VTA_DA_ neurons did not reduce consumption of foods with known caloric value (Fig. 1d). However, it did reduce the consumption of food with a larger than expected caloric content (i.e. on the first day that a flavour was paired with closed-loop nutrient infusion; Fig. 1c). This suggests that VTA_DA_ neurons are necessary for updating food preferences and consumption in response to new information about a food’s caloric content.

**Figure 1.**
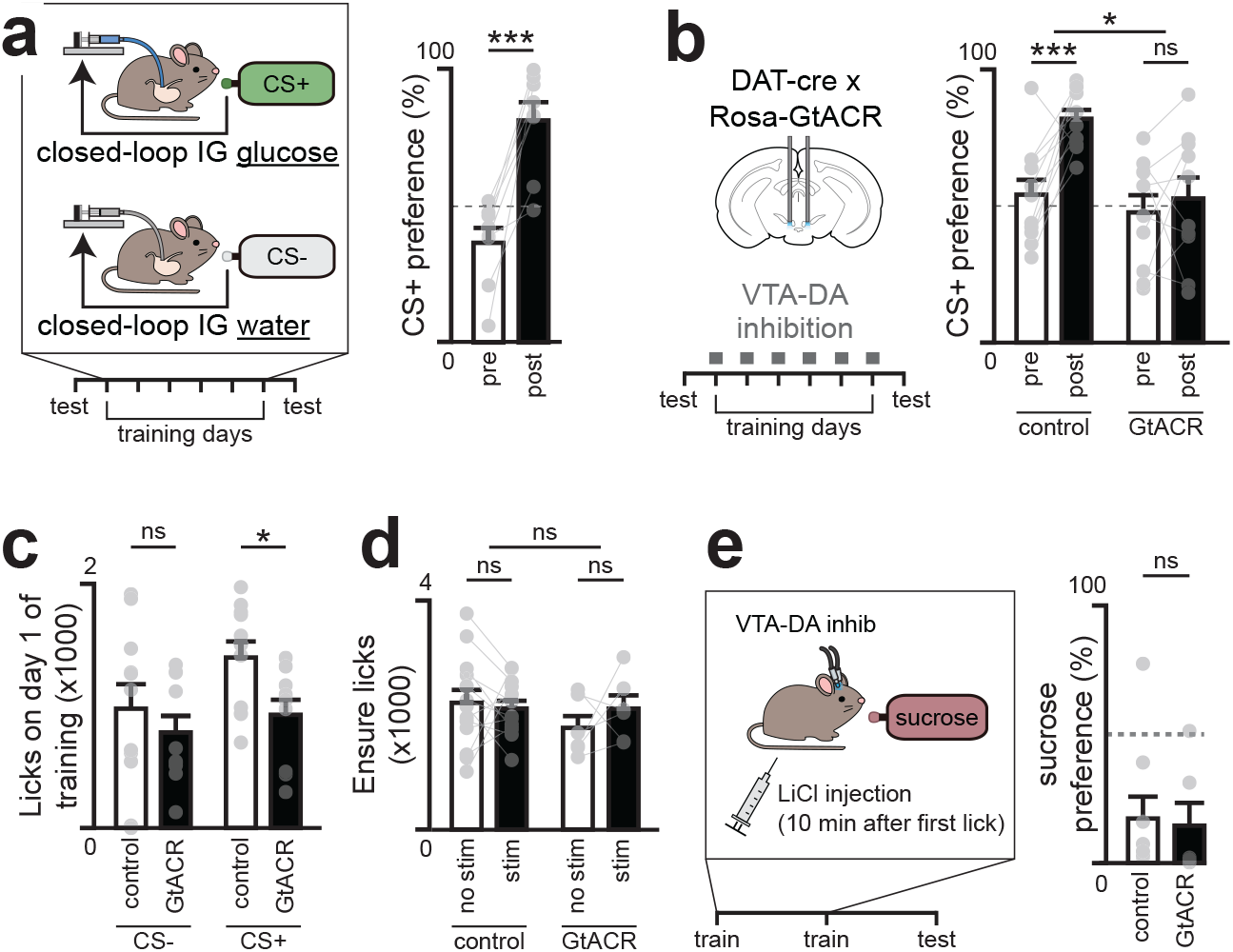
Dopamine neurons are necessary for nutrient learning. a, Flavour–nutrient conditioning protocol (left); mice learned to prefer the glucose-paired flavour (CS+, right). b, Inhibition of VTA-DA neurons throughout the entirety of the training session (left) prevented flavour–nutrient learning (right). c, VTA-DA inhibition reduced CS+ intake on the first training day. d, Inhibition did not reduce intake of the familiar nutritive solution Ensure. e, Flavour avoidance conditioning protocol (left); VTA-DA inhibition did not prevent acquisition of conditioned flavour avoidance (right). ns P > 0.05; *P < 0.05; **P < 0.01; ***P < 0.001. Error bars in bar plots show SEM. See Supplementary Table 1 for statistical tests, n, and p-values.

In addition to learning to prefer foods that have calories, animals also learn to avoid foods that are toxic, which occurs through an analogous post-ingestive learning process known as conditioned flavour avoidance^13–15^ (CFA). To test whether VTA_DA_ neurons are involved in learning about toxins, we asked whether silencing these cells could block the CFA that results from pairing a novel flavour with LiCl injection, which causes visceral malaise (Fig. 1e). In contrast to flavour-nutrient learning (Fig. 1b), we found that silencing VTA neurons had no effect on flavour-toxin learning (Fig. 1e). This indicates that VTA_DA_ neurons are specifically required for learning about nutrients but are not necessary for all forms of post-ingestive learning.

### Nutrients selectively trigger DA release in the BLA

We hypothesized that DA neurons contribute to flavour nutrient conditioning by supplying a signal of nutrient detection from the gut. Of note, previous studies using microdialysis have reported nutrient triggered DA release in striatal sites (such the nucleus accumbens core, NAc, and the dorsolateral striatum, DLS^16–24)^ and this is assumed to represent post-ingestive reward. However, we recently observed dramatic post-ingestive DA release in basolateral amygdala (BLA)^25^. We therefore systematically compared DA release across these brain regions.

We targeted GRAB-DA3m^26^ to NAc, DLS, and two sites in the BLA (medial, mBLA; and lateral, latBLA) for fiber photometry recordings of extracellular DA and in the same surgery installed IG catheters for nutrient infusion (Fig. 2a; Extended Fig. 2). Infusion of the complete diet Ensure caused dramatic DA release in mBLA (18.2±3.5%, p=1e-5) which began 48±11 seconds after nutrients entered the stomach and remained elevated for 9±1 minutes after the infusion ended (Extended Fig. 2af-n). This response was specific to nutrients, as it was also observed for infusion of glucose but not water (13.5±4.0% vs. -0.1±0.3% p=3e-5) or mannitol (Extended Fig. 2i), and it was independent of sex, as it was observed in both male and female mice (Extended Fig. 2h). In contrast, less nutrient-triggered DA release was observed in latBLA and none in dorsal or ventral striatum (Fig. 2a; Extended Fig. 2o-v). This is consistent with our prior finding that nutrient detection in the gut does not trigger DA release in the other major targets of the mesolimbic DA pathway^25^.

**Figure 2.**
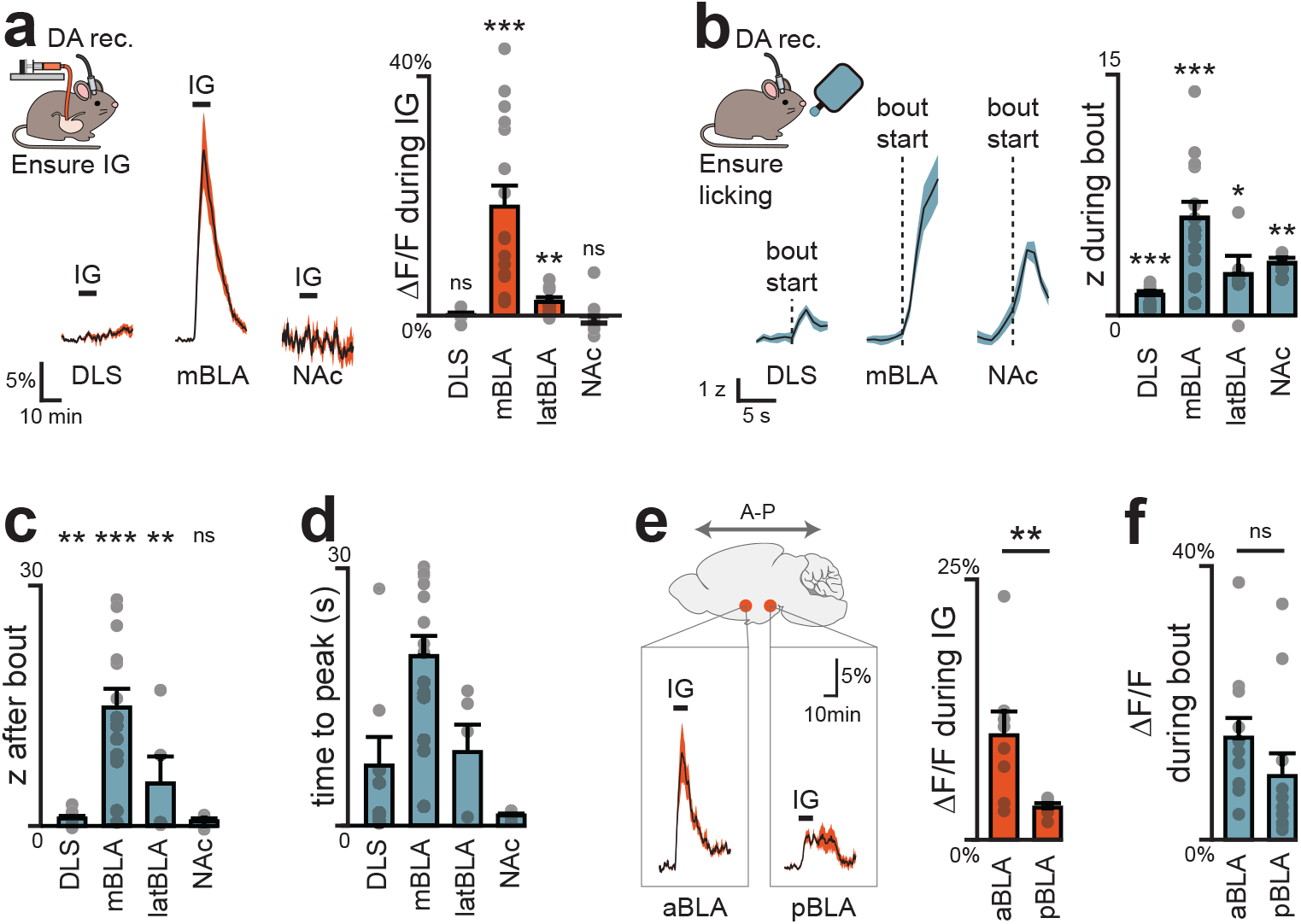
Distinct dopamine pathway tracks internal nutrients. a, Dopamine release recorded with the fluorescent sensor GRAB-DA during Ensure IG infusion, showing responses localized to BLA. b, Dopamine release 5 s into the first lick bout, showing distributed responses. c, Dopamine release 5 s after the first lick bout, showing sustained activity in BLA. d, Peak times within 30 s of the first lick bout. Note prolonged BLA responses. e, During IG infusion, dopamine release was greatest in anterior BLA (aBLA). f, During the first lick bout, dopamine release was similar across BLA subregions. Mice were sated for all infusion recordings (here and elsewhere), unless otherwise stated. ns P > 0.05; *P < 0.05; **P < 0.01; ***P < 0.001. Shaded areas in mean traces and error bars in barplots show SEM. See Supplementary Table 2 for statistical tests, n, and p-values.

To confirm that these DA dynamics in BLA also occur during natural ingestion, we next recorded DA release during self-paced consumption of Ensure (Fig. 2b; Extended Fig. 3). In mBLA, we observed a dramatic increase in DA release that ramped during the first minute of ingestion and was sustained for the entire trial (Fig. 2b–d; Extended Fig. 3a-e). In contrast, DA release in NAc peaked almost immediately after ingestion and then rapidly declined, consistent with known responses in NAc to appetitive tastes (Extended Fig. 3j-m). Interestingly, we observed a small but statistically significant (1.0±0.3%, p=1e-4; Extended Fig. 3q) increase in DA release in DLS after ingestion that may correspond to previous observations using microdialysis^18–21^. Together, this indicates that the mBLA is the major site of DA release in response to nutrient detection in the gut.

### DA release in response to nutrients is restricted to the anterior BLA

The BLA is well known for its role in assigning valence to exterosensory cues^27,28^. This has been most studied in the context of fear conditioning^29,30^, but the BLA is also involved in processing appetitive cues^27,28,31–34^. These effects have been attributed to different subregions of the BLA, which spans 2.5 mm in the rostro-caudal axis in the mouse with regional differences in cytoarchitecture and connectivity^35–38^. How interoceptive signals, such as post-ingestive nutrients, are represented across these subregions has not been examined. We observed the largest nutrient-triggered DA responses in the mBLA, which is part of the magnocellular region that extends to anterior BLA. Meanwhile the latBLA, which is part of the parvocellular region that extends posteriorly, showed markedly smaller responses (Extended Fig. 2s).

We therefore prepared mice for DA recordings in anterior vs. posterior BLA to test directly whether there is differential DA release across the rostro-caudal axis (Fig. 2e; Extended Fig. 4). This revealed that nutrient infusion caused much larger DA release in the anterior versus posterior BLA (e.g. Ensure: 10.0±2.3% vs. 3.1±0.4%, p=8.5e-3; Fig. 2e). Importantly, we found that the DA release in response to licking – which is immediate and does not involve GI signals – was the same in the anterior and posterior BLA, indicating that the difference in GI responses was not due to a technical defect in our posterior BLA recordings (Fig. 2f, Extended Figs. 4a, 4m-n). Indeed, we found that DA release in posterior BLA was preferentially triggered by aversive GI stimuli, such as injection of LiCl (Extended Fig. 4r) and LPS (Extended Fig. 4s), which did not cause DA release in anterior BLA. This shows that the anterior and posterior BLA respond to fundamentally different visceral signals.

We next investigated the gut sensors that detect nutrients upstream of the BLA. We reasoned that, if this pathway is involved in flavour-nutrient learning, then DA release in the anterior BLA should depend on the specific gut receptors that have been implicated in this process. For sugar, it has been shown that the key sensor is the intestinal sodium glucose transporter SGLT1^39,40^. Consistently, we found that IG infusion of a variety of SGLT1 agonists (glucose, galactose, and MDG) induced strong DA release in anterior BLA, whereas sugars that do not activate SGLT1 (mannitol and fructose) did not stimulate DA release (Extended Fig. 4g). Conversely, blocking intestinal SGLT1 with a specific antagonist was sufficient to prevent anterior BLA DA release in response IG glucose infusion (Extended Fig. 4h), indicating that this transporter is necessary. For fat, the key intestinal sensors are GPR40/120 and CD36^41–43^. Accordingly, we found that blocking all three of these intestinal receptors abolished DA release in the anterior BLA during fat infusion (Extended Fig. 4j). Thus, DA release in the anterior BLA depends on the same gut sensors that are known to drive flavour-nutrient conditioning.

### CCK+ DA neurons that project to BLA preferentially respond to nutrients

Our finding that nutrients trigger DA release only in the anterior BLA suggests that these responses are driven by only a small percentage of all VTA_DA_ neurons. Indeed, when we performed single-cell calcium imaging of all VTA_DA_ neurons, we observed that only 11% of cells were activated during IG nutrient infusion^25,44^ (Fig. 3a). We therefore set out to identify these nutrient-responsive DA neurons.

**Figure 3.**
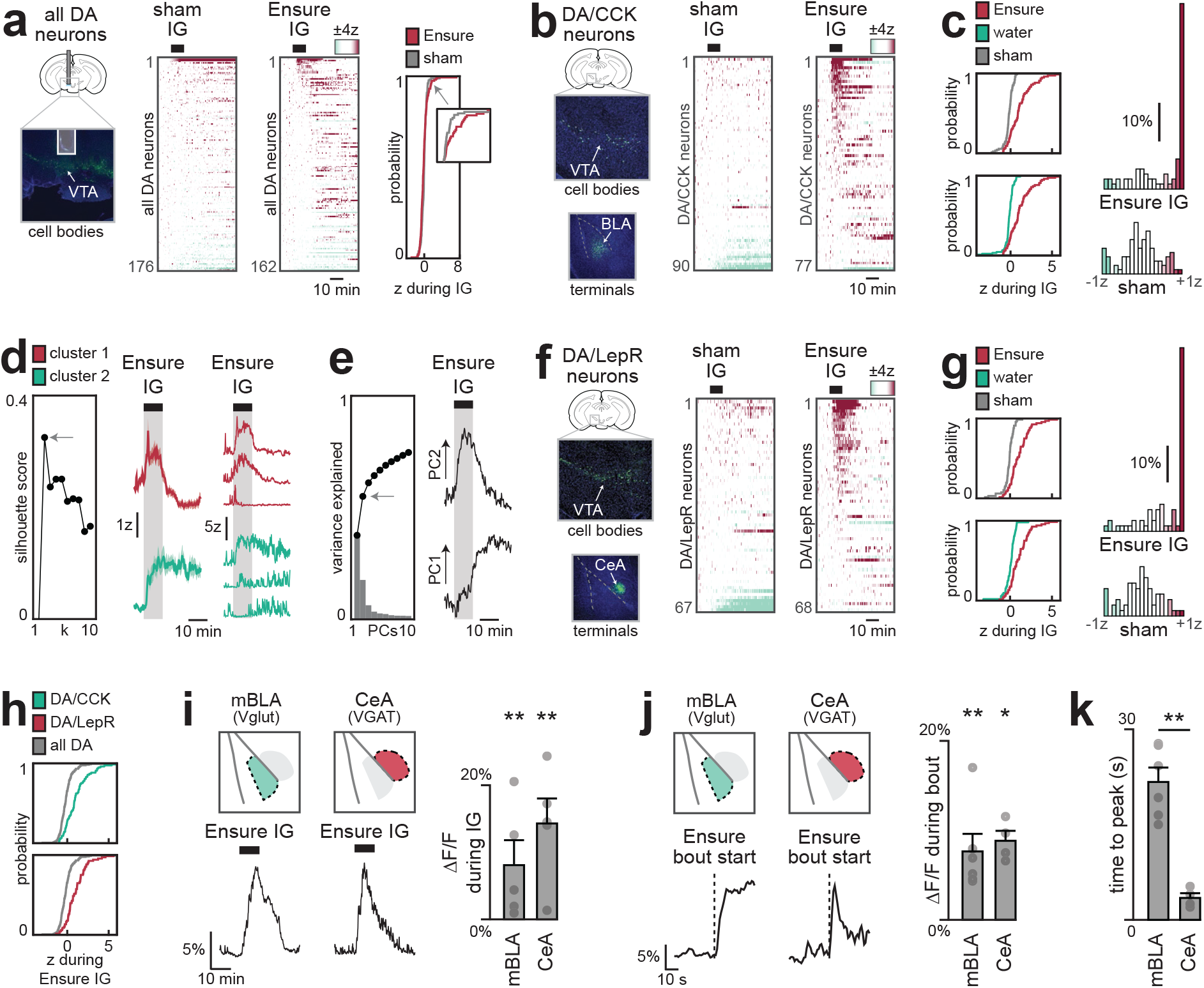
Genetically defined VTA dopamine projections track nutrient ingestion. a, Single-cell (miniscope) dynamics of VTA-DA neurons during Ensure or sham infusion. A small subpopulation showed greater activation during Ensure infusion vs sham (p = 0.018). b–e, VTA-DA/CCK neurons (cell bodies in VTA, terminals in BLA). b, Single-cell (miniscope) dynamics during Ensure or sham infusion. c, Cumulative distributions and binned activity show increased responses during Ensure IG vs sham (p = 2.0e-13) or water (p = 8.8e-13). d, K-means clustering reveals transient (magenta) and prolonged (mint) subpopulations, shown with mean and representative single-neuron activity. e, Principal component analysis; the first two PCs explain 55% of variance and correspond to activity changes during and after infusion. f–h, VTA-DA/LepR neurons (cell bodies in VTA, terminals in CeA). f, Single-cell (miniscope) dynamics during Ensure or sham infusion. Note lack of sustained Ensure response, relative to CCK subpopulation. g, Cumulative distributions and binned activity show increased responses during Ensure IG vs sham (p = 6.8e-10) or water (p = 4.9e-7). h, Cumulative distributions show increased activation across both CCK and LepR subpopulations (CCK, p = 4.2e-11; LepR, p = 2.3e-12). i–k, Fiber photometry recordings of DA release in mBLA and CeA using region-specific Cre drivers (Vglut-cre, VGAT-cre). i, Dopamine increased in both regions during Ensure IG (left: example traces; right: summary plots). j, Dopamine increased in both regions 5s into the first Ensure lick bout (left: example traces; right: summary plots). k, Lick-triggered DA release peaks earlier in CeA. Mice were sated for all infusion recordings (here and elsewhere), unless otherwise stated. ns P > 0.05; *P < 0.05; **P < 0.01; ***P < 0.001. Shaded areas in mean traces and error bars in bar plots show SEM. See Supplementary Tables 2 and 3 for statistical tests, n, and p-values.

Previous anatomic studies have shown that VTA_DA_ neurons expressing cholecystokinin (VTA_DA,CCK_) project preferentially to the BLA^45^, whereas VTA_DA_ neurons expressing the leptin receptor (VTA_DA,LepR_) project to the adjacent central amygdala^46^ (CeA). We therefore targeted GCaMP7s to these cells using an intersectional genetic approach and performed single-cell imaging in response to nutrient infusions (Fig. 3b; Extended Fig. 5). Consistent with our photometry results, we found that a majority of VTA_DA-CCK_ cells (61%) were activated during nutrient infusion in manner that mirrored the DA dynamics in the BLA (Fig. 3c). For example, somatic calcium ramped in the first minute after the onset of infusion (rise time: 36±7s for calcium vs. 41±12s for DA in BLA; see Methods; Extended Figs. 2k–l, 5e) and then remained elevated for many minutes after nutrient infusion ended (Extended Fig. 5g).

While most VTA_DA-CCK_ neurons responded to IG nutrients, these responses were not homogeneous. Indeed, k-means clustering revealed two populations of VTA_DA-CCK_ cells, one that was activated primarily during IG infusion and a second population of cells that showed more sustained responses (Fig. 3d). Consistently, principal component analysis (PCA) revealed two distinct phases of activity—one during nutrient infusion (PC2) and one after (PC1) (Fig. 3e)— and this first principal component was sufficient to segregate the two clusters of cells identified by k-means (Extended Fig. 5f). This reveals that internal signals of nutrient ingestion – which persist for tens for minutes – are represented by two subpopulations of VTA_DA-CCK_ cells with distinct dynamics.

We found that VTA_DA-LepR_ neurons also showed strong and sustained responses to IG nutrient infusion (Fig. 3f–h; Extended Fig. 5p-y). Consistent with this, we found that IG nutrients triggered robust DA release in the CeA (Fig. 3i; Extended Fig. 6). We confirmed that this DA release was distinct from the DA release in adjacent BLA by using region-specific Cre drivers (Vgat for CeA, Vglut2 for BLA) to confine GRAB-DA to the respective nuclei. Moreover, we found that the DA release during natural ingestion peaked much earlier in CeA compared to BLA (3.4±0.7 s vs 21.7±2.3 s, p=4.7e-3; Fig. 3j–k; Extended Figs. 6c–d, 6h–i), suggesting that DA release in these two structures may have distinct functions.

### Only the dopamine pathway to anterior BLA drives flavour preference

We next investigated which of these DA pathways is involved in learning about nutrients and flavours (Fig. 4a). We prepared mice for optogenetic stimulation of VTA_DA_ terminals in the anterior BLA and CeA, which both show strong nutrient-induced DA release, as well three other control sites: the posterior BLA, which we found responded to aversive visceral signals (Fig. 2,3, Extended Figs. 2–4, 6), and the NAc and DLS, which are important for many other forms of DA-mediated learning^47,48^.

**Figure 4.**
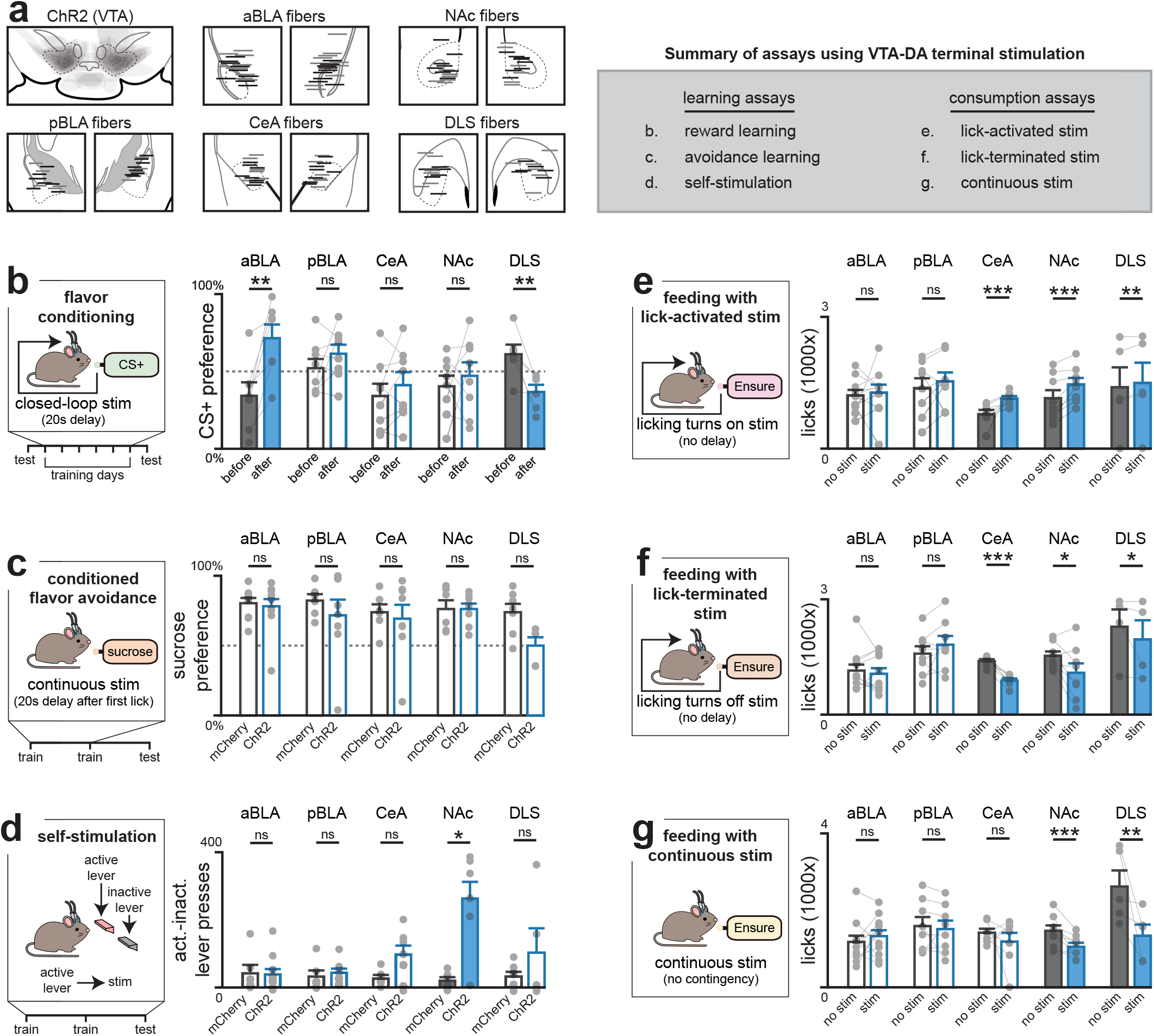
Distinct roles of BLA dopamine projections in feeding and flavour learning. a, (Top) Viral expression of ChR2 in VTA-DA neurons and fiber placement above projection targets (grey=mCherry, black=ChR2, left) and summary of assays (right). b, Flavour conditioning assay (during training, each lick triggered stimulation after a 20s delay, replicating IG transit time). c, Conditioned flavour avoidance assay (during training, continuous stimulation was given after the first lick with a 20s delay). d, Self-stimulation assay (during training, the active lever provided stimulation with no delay). e, Consumption during lick-activated stimulation (no delay). f, Consumption during lick-terminated stimulation (no delay). g, Consumption during continuous stimulation (no contingency). ns P > 0.05; *P < 0.05; **P < 0.01; ***P < 0.001, after Benjamini–Hochberg correction. Error bars in bar plots show SEM. See Supplementary Table 1 for statistical tests, n, and p-values.

We reasoned that if nutrient-triggered DA release in a brain region drives flavour learning, then optogenetically-induced DA release in that structure should substitute for IG infusion. Mice were given access to two differently flavoured, non-caloric solutions, one of which was paired with stimulation following a 20s delay (to mimic the time course of post-ingestive DA release; see Methods and Extended Data Fig. 2j), while the other solution did not trigger stimulation. Strikingly, this delayed stimulation of DA release in anterior BLA conditioned a robust flavour preference (Fig. 4b; Extended Fig. 7a), whereas no increase preference was observed after stimulation of axon terminals in any other region. This indicates that only the anterior BLA projection is sufficient to drive flavour learning.

To evaluate the specificity of these results, we tested these mice in a battery of assays measuring different aspects of feeding behavior or reinforcement. In contrast to the effects on flavour learning, stimulation of axon terminals in the anterior BLA had no effect on total food intake, regardless of the timing of optogenetic stimulation (either continuous or closed-loop to lick onset or offset; see Methods; Fig. 4e–g; Extended Fig. 7b-d) and also failed to condition avoidance of a paired flavour (Fig. 4c). This pathway also failed to support self-stimulation, indicating it is not reinforcing in the absence of a flavour cue^49^ (Fig. 4d). Thus, DA release in anterior BLA influences feeding by specifically driving flavour learning.

Although we saw similar dynamics of nutrient-triggered DA release in CeA, stimulation of the VTA to CeA pathway did not drive flavour learning (Fig. 4b). However, it did potentiate ongoing feeding and drinking behavior, consistent with the idea that DA release in this structure is reinforcing (Fig. 4e–g; Extended Fig. 7b-f). We observed similar effects when stimulating the pathway to NAc^50,51^ (Fig. 4e–g; Extended Fig. 7b-f). On the other hand, DA release in DLS caused a reduction in flavour preference (61.1±6.2% before vs. 36.7%±4.6% after training; p=1.1e-3, Fig. 4b) and there was a trend toward conditioned flavour avoidance (73.9±6.0% vs. 50.0±6.0% sugar preference; p=1.3e-2; Fig. 4c). Taken together, these results reveal a double dissociation in which only the anterior BLA pathway drives flavour learning in response to delayed stimulation, while other aspects of motivated behavior or reinforcement are mediated by canonical DA pathways.

### Anterior BLA neurons integrate gustatory and post-ingestive signals to drive learning

The fact that nutrients trigger DA release in anterior BLA, and this drives flavour learning, raises the question of how flavours and post-ingestive signals are integrated in this structure. In pilot studies, we found that IG nutrient infusion was unable to condition a preference when it was separated in time from the paired flavour (Extended Fig. 8a), implying that these signals must overlap for learning to occur (which is in contrast to CFA, which permits long delays between flavour and toxin^14^). We therefore prepared mice for two-photon imaging of anterior BLA neurons expressing dopamine receptor 1 (D1R), which labels most DA-responsive BLA cells^52^ (Fig. 5a), and examined how these cells integrate ongoing nutrient and flavour signals.

**Figure 5.**
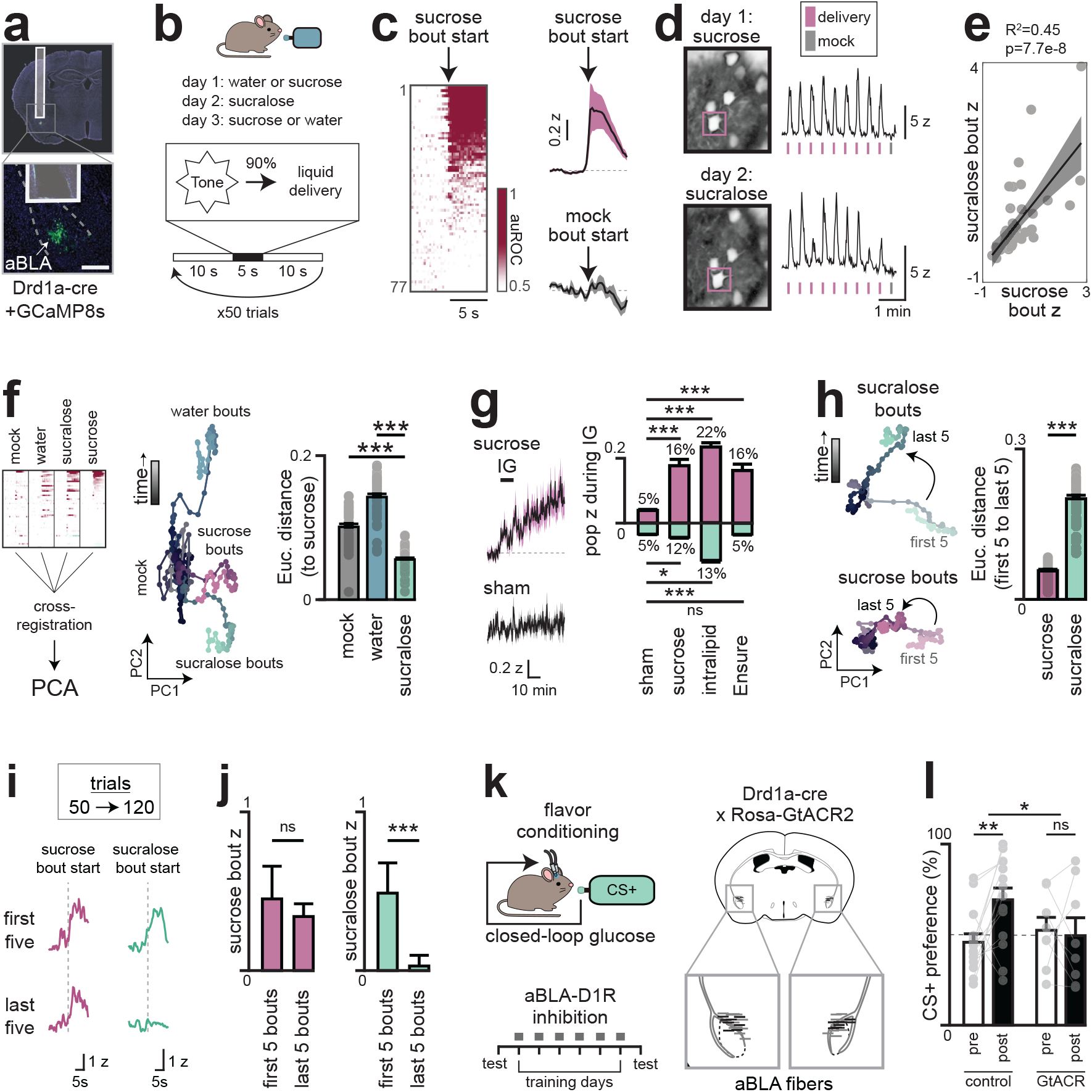
aBLA neurons integrate oral and post-ingestive nutrient signals. a, Imaging of D1R-expressing aBLA neurons (scale bar, 0.5 mm, 2-photon). b, Protocol for recording taste responses. c, Single-cell dynamics during sucrose delivery, quantified by auROC (left), and mean activity across sucrose vs no-delivery bouts (right). d–e, Cross-registered neurons across days. d, Example FOVs and activity of a single neuron during sucrose (day 1) and sucralose (day 2) delivery. e, Correlation of activity during sucrose vs sucralose bouts. f, PCA of cross-registered neurons across three days shows similar activity trajectories for sucrose and sucralose, quantified by euclidian distance across PC1–2 for each timepoint. g, Mean activity traces during IG infusions (left), and population-weighted z-scores of activated (magenta) and inhibited (mint) neurons (right). h, Lick-triggered activity diverged across early and late sucralose (but not sucrose) bouts, shown in PC space (left) and by Euclidean distance across PC1–2 (right). i–j, Increasing trials to 120 reveals further changes in responsivity. i, Activity of one neuron during early and late sucrose (left) and sucralose (right) bouts. j, Sucralose (but not sucrose) responses decayed over time. k–l, Inhibition of aBLA-D1R neurons during flavour–nutrient conditioning. k, Protocol (left) and fiber placement (right; grey=littermate controls, black=GtACR). l, Silencing throughout the entirety of the training session prevented conditioned flavour preference. Mice were sated for all infusion recordings (here and elsewhere), unless otherwise stated. ns P > 0.05; *P < 0.05; **P < 0.01; ***P < 0.001. Shaded areas in mean traces and error bars in bar plots show SEM. See Supplementary Tables 1 and 3 for statistical tests, n, and p-values.

We first sought to isolate BLA responses to flavour cues by recording activity during brief-access taste tests (5 seconds, each trial initiated by an auditory cue) for sucrose, sucralose (a non-caloric sweetener), or water. To distinguish taste-driven activity from other cue responses^38^ or licking itself, in 10% of trials (mock trials) the cue was played but liquid access was withheld (Fig. 5b).

We found that 58% of anterior BLA_D1R_ neurons were activated by sucrose ingestion with dynamics that were time-locked to the start of licking (Fig. 5c; peak time: 3.4±0.3s from the first lick in each bout; Extended Fig. 8b-c). In contrast, we observed only weak activation of these cells during mock trials (1.0±0.2z vs. 0.08±0.09z, sucrose vs. mock, p=1e-5), and these differences persisted when we controlled for the number of licks (Extended Fig. 8d-f), indicating these responses depend on ingestion. In addition, the responses to sucrose and sucralose were highly correlated, both at the level of individual neurons (p=7.7e-8; Fig. 5d–e; Extended Fig. 8g– i) and population dynamics (Fig. 5f; Extended Fig. 8j). In contrast, ingestion of water produced less activation and a distinct population trajectory (Fig. 5f; Extended Fig. 8h-j). Together, this demonstrates that many anterior BLA neurons specifically represent sweet tastes.

We next investigated how anterior BLA_D1R_ neurons respond to purely post-ingestive signals by recording their activity during and after IG infusion of sucrose. This revealed a slow ramping activation during sucrose infusion that was not observed with sham, water or non-nutritive mannitol infusion (Fig. 5g; Extended Fig. 8k-p). We observed a similar ramping activity following infusion of Intralipid or Ensure, confirming that these are bona fide nutrient responses.

We wondered how the phasic signal representing sweet taste and the tonic signal represent GI feedback are integrated in individual neurons. We reasoned that if learning happens in the anterior BLA, then nutrients might modulate the strength of BLA responses to gustatory signals Indeed, we found that responses to licking non-nutritive sucralose diminished during the course of the one session, whereas responses to sucrose were stable at the single-cell and population level (Fig. 5h–j; Extended Fig. 8q-r). This indicates that nutrient and gustatory responses converge on the same cells in anterior BLA (similarly to Insula^53^), and that this integration strengthens responses to tastes associated with calories.

The integration of taste and post-ingestive signals in anterior BLA_D1R_ neurons suggests that these cells could be critical for learning about the nutrient content of food. To test their necessity, we optogenetically silenced anterior BLA_D1R_ neurons during flavour-nutrient conditioning, which completely prevented mice from learning which flavour was associated with calories (Fig. 5k–l; Extended Fig. 9a). Importantly, silencing these neurons did not result in a decrease in food intake when hungry animals were given Ensure (Extended Fig. 9c), indicating that anterior BLA_D1R_ neurons are not required for ingestion per se. However, silencing these neurons did decrease consumption of a non-caloric solution newly paired with IG glucose infusion (Extended Fig. 9b), indicating that they are important for updating the value of flavours when the nutrient content of food is uncertain. This effect was specific to flavour-nutrient learning, as inhibition of these neurons did not block a different form of post-ingestive learning, CFA (Extended Fig. 9d). Thus, anterior BLA_D1R_ neurons integrate flavour cues with nutrient signals to assign value to calorie-rich foods.

### Post-ingestive nutrients preferentially engage the left BLA

All the recordings described above were performed in the left amygdala. However, there have been reports that aversive stimuli, such as those that induce fear^54^ and pain^55^, preferentially engage the right amygdala. We therefore compared responses in the right and left BLA during nutrient infusions. This revealed that the minutes-timescale response to IG nutrients was greater in left BLA (18.2±3.5% vs 5.3±1.0%, p=5.3e-4; Extended Figs. 10a, 11a-c) whereas the seconds-timescale lick-triggered response was larger in the right (Extended Figs. 10b, 11d–f). We confirmed this result by recording simultaneously from both hemispheres in a new cohort of mice (Extended Figs. 10c–d, 11g–k). This demonstrates that ingested nutrients trigger dopamine release preferentially in the left BLA in mice.

We wondered whether the left BLA was also engaged in humans by post-ingestive nutrients. A previous study showed that dopamine release occurred within the BLA after ingestion of a milkshake^56^, but this study was not powered to measure absolute differences across hemispheres (n=10 subjects; see Methods; Extended Figs. 10e, 11l–m). We therefore designed an fMRI experiment (n=38 subjects) to test the hypothesis that BLA activity is lateralized during the post-ingestive period in humans. Participants consumed either a glucose or water solution, and BOLD signals were measured during and after ingestion. This revealed post-ingestive activity was greater in the left BLA than in the right (p=0.005; Extended Figs. 10f, 11n; Supplementary Table 4), indicating that nutrient signals are routed to the left BLA in both mice and humans.

We considered two possible mechanisms that could give rise to these lateralized responses (Extended Fig. 10g). One possibility is that they arise from asymmetric sensory inputs to the brain. This is suggested by data showing that there are functional differences between the left and right vagus nerve^19,57^. We tested this by removing the vagus from one side of the body and measuring DA responses in the left and right BLA. This revealed that removing either vagus reduced nutrient responses in both hemispheres (Extended Figs. 10h–i, 11o–z), ruling out the hypothesis that lateralization arises from separate asymmetric pathways.

An alternative possibility is that lateralization is imposed by a top-down pathway, suggested by the fact that insular cortex, which is heavily interconnected with BLA, shows asymmetric responses to aversive visceral cues^58,59^. To test whether there are differences in the inputs to the right versus left VTA^DA^—>BLA pathway (Extended Fig. 12a-e), we injected a Cre-dependent pseudorabies virus (PRV-Introvert-mCherry) into either the left or right BLA of DAT-Cre mice (Extended Fig. 10j), which spreads retrogradely at a rate of approximately one synapse per day, and then performed brain-wide imaging and neuron registration four days later. Strikingly, we found that the most highly enriched structure was visceral cortex, a subdivision of the insula known to be functionally lateralized^58,59^ and the next top 10 regions included several interconnected higher-processing structures, (Extended Figs. 10k–m, 12f–l). This prioritizes further investigation of the insula-amgydala network to understand how lateralization arises.

## Summary

Animals must learn which foods are nutritious based on their post-ingestive effects^1,2^. While these post-ingestive effects could in principle be associated with any sensory cue, there is much evidence that there is a privileged pathway connecting the flavour of food, sensed in the mouth, with its nutrient content sensed in the gut. For example, animals can readily learn to lick a flavoured solution, but not press a lever, to receive an IG infusion of nutrients^3,4^. Similarly, animals will learn to prefer a place associated with IG nutrient infusion only if it is coupled with an orosensory cue^4^. This suggests that there is a hardwired pathway in the brain that connects flavours and nutrients to drive learning about food, but the underlying circuit-level mechanisms have remained unclear.

Here, we have shown that there is a genetically and anatomically-distinct pathway within the midbrain dopamine system that is responsible for associating nutrients and flavours. This pathway is defined by CCK-expressing VTA_DA_ neurons that relay internal nutrient signals to the anterior portion of the BLA. A variety of functional tests show that this pathway receives the specific GI signals known to drive flavour preference, and that the underlying neurons are necessary and sufficient for flavour learning. On the other hand, we show that striatal DA pathways, which have long been assumed to drive this process^16–20,22–24^ show weak or absent responses to GI nutrients and are not sufficient to drive flavour learning. This identifies a dopaminergic circuit involving the BLA—which has previously been shown to rapidly respond to motivationally salient food cues^38^ and encode reward identities^49^—to be critical for using delayed internal signals to learn which foods are nutritious.

Learning about food involves making associations with both its nutrient content and its environmental context. The latter is known to be mediated by phasic DA release in NAc in response to reward-predicting exterosensory cues. A salient feature of this form of learning is that it is temporally precise, meaning that DA must be released within (at most) a few seconds of the cue^60^. In contrast, nutrient detection in the gut is inherently slow and, as we show, involves DA signals that ramp over tens of minutes in the BLA. Consistent with the distinct properties of these two pathways, we found that acute stimulation of NAc DA projections invigorated ongoing behavior, but delayed and prolonged stimulation did not drive flavour learning. Conversely, acute stimulation of BLA projections did not alter behavior, but delayed stimulation drove robust flavour learning. This suggests that the BLA pathway drives (or permits^49^) flavour–nutrient associations, while other pathways (e.g., NAc) link those flavours to external cues. An important task for the future will be to elucidate the molecular and synaptic mechanisms that differentiate these two pathways and enable learning on such different timescales.

## Methods

All experimental protocols using mice were approved by the University of California, San Francisco Institutional Animal Care and Use Committee, in accordance with the NIH Guide for the Care and Use of Laboratory Animals. Human experiments were approved by the ethics committee of the Medical Faculty of the University of Cologne, Germany (fMRI, No. 17-278; PET, No. 16-320). For clarity, methods are organized into sections for surgery, recordings, behavior, and analysis. Each subsection specifies the species used (mouse or human).

### Mouse strains

The following transgenic lines were used: Drd1a-Cre (MMRRC #030989), DAT-Cre (Jackson #006660), CCK-Cre (Jackson #012706), Vglut2-Cre (Jackson #016963), Vgat-Cre (Jackson #016962), DAT-Flp (Jackson #033673), TiGRE-GCaMP7s (Jackson #034112), Rosa-GtACR1 (Jackson #033089), and RosaFL-ReaChR (Jackson #024846). LepR-Cre mice were obtained from Martin Myers. Double and triple mutant lines were generated through standard breeding. Mice used for experiments ranged from 6 weeks to 9 months old, and were represented in roughly equal number of males and females.

### SURGERY: General methods (mice)

Mice were anesthetized with 2% isoflurane and placed on a thermostatically controlled heating pad. Sustained-release buprenorphine (1.5 mg/kg, subcutaneous, or SC) and meloxicam (5 mg/kg, intraperitoneal, or IP) were administered preoperatively. Ophthalmic ointment was applied to the eyes, and surgical sites were shaved and sterilized with alternating betadine and ethanol. Local anesthesia was provided with 0.25% bupivacaine (SC) at the incision site.

### SURGERY: Intragastric catheters (mice)

Sterile polyurethane catheters (C30PU-RGA1439, Instech Labs) were attached to vascular access buttons (VABM1B/22, Instech Labs). Catheters were inserted into the avascular wall of the stomach and externalized dorsally through the access button secured on the upper back. Mice recovered for at least one week before experimentation.

### SURGERY: Unilateral vagotomies (mice)

Under isoflurane anesthesia, a midline ventral neck incision was made to expose the salivary glands, which were gently separated and retracted laterally. The sternomastoid, sternothyroid, and omohyoid muscles were then retracted to reveal the carotid triangle. The carotid sheath was carefully opened, and the vagus nerve isolated for transection (>2 mm segment removed) below the level of the superior laryngeal branch. The incision was immediately closed with absorbable sutures, and mice were allowed to recover. In rare cases, transient ipsilateral ptosis (Horner’s sign) was observed, likely due to inflammation of the cervical sympathetic trunk; otherwise, the procedure was well tolerated without complications.

All recordings were performed within two weeks after surgery, and post-mortem inspection confirmed no visible nerve regrowth. In a subset of mice, intraperitoneal injection of WGA Alexa Fluor 555 was performed after experimental completion. One week later (3 weeks after surgery), nodose ganglia were dissected and examined, revealing labeling restricted to the ganglion contralateral to the vagotomy, confirming lasting nerve transection.

### SURGERY: Intracranial injections (mice)

Mice were anesthetized with isoflurane and positioned in a stereotaxic apparatus (Kopf Instruments). The skull was leveled between bregma and lambda, and small burr holes were drilled above target regions. Viral vectors were delivered via pulled glass pipettes. Pipettes were left in place for >5 min after injection to prevent backflow, then raised 0.05 mm dorsoventrally to create slight negative pressure and held for an additional >5 min before withdrawal. Coordinates are given relative to bregma.

#### Anterograde tracing

(see Fig. 3b) CCK-Cre × DAT-Flp mice were prepared for tracing VTA-DA/CCK terminals by injecting AAV9-Con/Fon-ChR2-GFP (300 nL; 6.0 × 10¹² vg/mL; provided by K. Yackle and I. Bachmutsky) bilaterally into VTA (−3.1 AP, ±0.5 ML, −4.5 DV). Mice were perfused three weeks later and their brains processed for histology. Direct injection into VTA was necessary here due to the presence of DA/CCK neurons in dorsal raphe and the periaqueductal grey, which project to CeA^45^. Note that tracing of VTA-DA/LepR terminals (see Fig. 3f) was accomplished instead with a genetic cross LepR-Cre × DAT-Flp with RosaFL-ReaChR mice.

#### Rabies tracing

(see Extended Fig. 12a-e) DAT-Cre mice were prepared for monosynaptic retrograde tracing by injecting AAV5-EF1α-FLEX-TVA-mCherry (400 nL; 4.9 × 10¹² vg/mL; UNC Vector Core) bilaterally into VTA (−3.1 AP, ±0.5 ML, −4.5 DV). Three weeks later, CVS-N2c-ΔG-mGFP (100 nL; 7.8 × 10⁸ vg/mL; Janelia) was injected into the left anterior BLA (−1.0 AP, −3.0 ML, −4.8 DV). Mice were perfused one week later and brains processed for histology.

#### Pseudorabies tracing

(see Extended Figs. 10j–m; 12f–l) DAT-Cre mice received PRV-Introvert-GFP (150 nL; 4.7 × 10⁹ vg/mL; Princeton) into either the left or right BLA (−1.7 AP, ±2.9 ML, −4.8 DV) for polysynaptic retrograde tracing. Mice were perfused 1–5 days after injection. Labeling first appeared in ipsilateral VTA (day 1), expanded within VTA by day 2, and reached bilateral hypothalamic nuclei (LH, PVH) by day 3. On day 4, labeling bilaterally extended to cortical, amygdalar, and hindbrain regions. By day 5, animals displayed systemic illness, and labeling had spread to both nodose ganglia. All analyses used brains collected 4 days post-injection—the latest stage showing extensive labeling with minimal distress.

### SURGERY: Intracranial implants (mice)

Following viral injection, optical fibers or GRIN lenses were implanted 0.1–0.15 mm above the target site and secured with dental cement (Metabond). A custom titanium headplate (eMachineShop) was attached with the GRIN lens to accommodate our 2-photon imaging set up. For miniscope imaging, a second surgery was performed 4 weeks after virus injection to install a baseplate (Inscopix 100-000279) above the GRIN lens. The baseplate was affixed with dental cement and protected with a removable cover (Inscopix 100-000241).

#### Dopamine imaging

(see Figs. 2, 3, and Extended Figs. 2–4, 6, 10–11) Wild-type mice were injected with AAV9-hSyn-GRAB-DA3m (200 nL; 4–8 × 10¹² vg/mL; WZ Biosciences) and implanted with an optical fiber (Doric MFC_400/430-0.48_6 mm_MF2.5_FLT) at the following injection coordinates:

- Left DLS (+1.5 AP, -1.5 ML, −3.0 DV)
- Left NAc core (+1.4 AP, −1.0 ML, −4.5 DV)
- Left or right BLA (−1.7 AP, ±2.9 ML, −4.8 DV)
- Left lateral BLA (−1.7 AP, −3.2 ML, −4.8 DV)
- Left anterior BLA (−1.0 AP, −3.0 ML, −4.8 DV)
- Left posterior BLA (−2.4 AP, −3.3 ML, −4.7 DV)

Simultaneous bilateral recordings used dual injections and fiber implants in the same animal.

For region-specific recordings (see Fig. 3i–k and Extended Fig. 6), Vglut2-Cre mice (BLA) and Vgat-Cre mice (CeA) were injected with AAV9-hSyn-DIO-GRAB-DA3m (200 nL; 8.8 × 10¹² vg/mL; WZ Biosciences) and implanted with fibers above left BLA (−1.7 AP, −2.9 ML, −4.8 DV) or left CeA (−1.5 AP, −2.65 ML, −4.5 DV).

#### VTA-DA neuron imaging

(see Fig. 3a and Extended Fig. 5a-c) For VTA-DA population imaging, DAT-Cre mice received AAV1-FLEX-CAG-jGCaMP7s (300 nL; 6.3 × 10¹² vg/mL; Addgene 104495) into left VTA (−3.1 AP, −0.5 ML, −4.5 DV), followed by implantation of a GRIN lens (8.4 × 0.5 mm, Inscopix 1050-004629) positioned 0.15 mm above the target.

For subpopulation imaging (see Fig. 3b–h and Extended Figs. 5d–y), CCK-Cre × DAT-Flp × RosaFL-GCaMP7s mice (for VTA-DA/CCK) and LepR-Cre × DAT-Flp × RosaFL-GCaMP7s mice (for VTA-DA/LepR) underwent the same GRIN lens implantation above left VTA (−3.1 AP, ±0.5 ML, −4.5 DV).

#### BLA-D1R neuron imaging

(see Fig. 5a–j and Extended Fig. 8) For recording anterior BLA neurons, Drd1a-Cre mice were injected with AAV1-CAG-FLEX-GCaMP8s (200 nL; 5.9 × 10¹² vg/mL; HHMI Janelia) into left aBLA (−1.0 AP, −3.0 ML, −4.8 DV) and implanted with a GRIN lens (8.4 × 0.5 mm, Inscopix 1050-004629) 0.15 mm above the injection site.

#### VTA-DA projection stimulation

(see Fig. 4 and Extended Fig. 7) For optogenetic activation of VTA dopamine terminals, DAT-Cre mice were injected with AAV1-EF1α-DIO-hChR2(H134R)-mCherry (300 nL; 6.0 × 10¹² vg/mL; Addgene 20297) into left and right VTA (−3.1 AP, ±0.5 ML, −4.5 DV). For optogenetic activation of dopamine terminals in DLS, DAT-Cre mice were injected with AAV1-EF1α-DIO-hChR2(H134R)-mCherry (300 nL; 6.0 × 10¹² vg/mL; Addgene 20297) in both bilateral VTA (−3.1 AP, ±0.5 ML, −4.5 DV) and bilateral substantia nigra (−3.1 AP, ±1.5 ML, −4.5 DV).

Paired optical fibers (200 µm, Thorlabs CFLC230-10) were implanted bilaterally 0.1 mm above downstream projection targets, including NAc (+1.4 AP, ±1.0 ML, −4.5 DV), aBLA (−1.0 AP, ±3.0 ML, −4.8 DV), pBLA (−2.4 AP, ±3.3 ML, −4.7 DV), CeA (−1.5 AP, ±2.65 ML, −4.5 DV), and DLS (+1.5 mm AP, -1.5 mm ML, -3.0 mm DV). Control mice received AAV1-hSyn-DIO-mCherry (300 nL; 5.7 × 10¹² vg/mL; Addgene 50459) instead of ChR2.

#### VTA-DA and aBLA-D1R neuron inhibition

(see Figs. 1, 5 and Extended Fig. 1, 9) For VTA-DA inhibition, DAT-Cre × Rosa-GtACR mice (and littermate controls) were implanted with bilateral fibers (Doric DFC_200/245-0.37_5 mm_GS1.0_FLT) 0.1 mm above VTA (−3.1 AP, ±0.5 ML, −4.5 DV).

For aBLA-D1R inhibition, Drd1a-Cre × Rosa-GtACR mice (and controls) were implanted with bilateral 200 µm fibers (Thorlabs CFLC230-10) 0.1 mm above aBLA (−1.0 AP, ±3.0 ML, −4.8 DV).

### RECORDINGS: General methods (mice)

Except where noted, recordings were performed in sound-isolated behavioral chambers (Coulbourn Habitest Modular System). Chambers were cleaned between sessions to remove residual olfactory cues. Mice were habituated to the chamber for ≥1 h with recording equipment and intragastric (IG) catheters attached before the first experimental day, and for 10–20 min before each subsequent session. When used for imaging, the excitation LED was turned on during habituation to reduce bleaching artifacts. Except when noted, mice were sated (not food/water deprived).

For recordings of BLA neuron dynamics, mice were head-fixed using the custom titanium headplate (eMachineShop, see surgery methods above) above a miniature treadmill. These mice were habituated for at least three 1 hour sessions, then re-habituated for 10–20 min before each recording. Both sexes were used in approximately equal numbers.

Lick-evoked responses were recorded while mice consumed specific solutions after 18–24 hours of water or food deprivation. Water-restricted mice received water, 32% sucrose or 0.8% sucralose (matched for perceived sweetness to sucrose solution); food-restricted mice received 35% Ensure (isocaloric to sucrose solution). Each session began with a 10-min baseline period followed by access to a lickometer spout. Mice were first habituated to the setup with 1 h access to sucralose while attached to the camera.

For imaging of BLA neuron dynamics, mice were trained to lick 32% sucrose in response to a 5-s 2 kHz tone cue. After three training sessions, recordings were performed over three consecutive days, each presenting water, sucrose, or sucralose in counterbalanced order across mice. Animals were presented with 50 or 120 cue trials, separated by 20 seconds (lasting 1250 or 3000 seconds, respectively). During these recording sessions, 10% of cues omitted liquid delivery to isolate licking-only trials (these no-delivery trials are called “mock” trials).

IG-evoked responses were recorded by infusing solutions through previously implanted catheters at 100 µL/min for 10 min (Harvard Apparatus 70-2001 syringe pump). Tubing (AAD04119 Tygon, LS20/PNP3M Instech) was connected before habituation. Infusions began after a 10-min baseline.

For receptor-blockade experiments, antagonists were mixed into the infused solution: 600 µg/mL mizagliflozin (SGLT1), 50 µg/mL GW1100 (GPR40), 50 µg/mL AH7614 (GPR120), or 10 µg/mL sulfo-N-succinimidyl oleate (CD36). For CCK blockade, 1 mg/kg devazepide (in saline, 10 mL/kg IP) was administered 30 min prior to infusion.

Sugar infusions (sucrose, glucose, fructose, galactose, MDG, mannitol) were 32% w/v; sucralose was 0.8%. Other nutrient solutions were isocaloric to 32% sucrose: 35% Ensure, 32% collagen peptide, 14% intralipid, diet A (17.3% sucrose + 4.5% intralipid + 4.5% peptide), and diet B (6.5% sucrose + 8.5% intralipid + 6.4% peptide). Controls used 14% mineral oil or double-distilled water.

Hormone-evoked responses were elicited after a 10-min baseline by IP injection of saline (10 mL/kg) containing 2 mg/kg serotonin HCl, 10–30 µg/kg cholecystokinin octapeptide, 100 µg/kg peptide YY, 2 mg/kg ghrelin, 2 mg/kg glucagon, or 10 µg/kg amylin.

Visceral malaise was induced by IP injection of saline containing lithium chloride (100 mg/kg) or lipopolysaccharide (200 µg/kg) after the same baseline period.

### RECORDINGS: Photometry (mice)

Fluorescence signals were acquired using continuous 470 nm and 420 nm excitation (40 µW per LED; RZ10X LUX-I/O, TDT). Signals were sampled at 1.0173 kHz using Synapse (TDT) and exported via Browser (TDT). Raw traces were downsampled to 4 Hz in MATLAB for analysis.

### RECORDINGS: Miniscope videos (mice)

Miniscope recordings were acquired at 8 Hz (0.2 mW/mm², 455 nm LED, gain = 8, 2× spatial downsampling) using an nVista 3.0 system (Inscopix DA 2.0). Videos were pre-processed by spatial (2×) and temporal (2×) downsampling, bandpass filtering, and motion correction in Inscopix Data Processing 1.9. Sessions were excluded if motion exceeded the average neuron diameter after correction.

Cell activity was extracted using CNMF-E (v1.1.0) implemented in MATLAB (github.com/zhoupc/cnmfe). Segmented ROIs were manually refined to remove motion artifacts, duplicates, and over-segmented regions. Cells exhibiting bleaching artifacts fit by an exponential decay during baseline were excluded.

### RECORDINGS: 2-photon videos (mice only)

Two-photon imaging was performed at 20 Hz (512 × 512 pixels, 900 nm excitation; Chameleon Discovery laser, Coherent) with a resonant-galvo scanner (Ultima Investigator, Bruker PV 5.8). Videos were spatially (2×) and temporally (2×) downsampled, bandpass-filtered, and motion-corrected using Inscopix Data Processing 1.9, followed by additional correction and ROI extraction using Ezcalcium (v3.0)^61^ built on NoRMCorre^62^ and CaImAn^63^ toolboxes (https://github.com/porteralab/EZcalcium). ROIs were manually curated with CaImAn refinement and cross-registered across sessions using custom code based on the CaImAn ROI-matching module.

### RECORDINGS: fMRI imaging (humans)

To test whether glucose consumption modulates BOLD activity in the human basolateral amygdala (BLA), overnight-fasted participants performed a Drink Task during fMRI (MAGNETOM Prisma Fit, Siemens AG, Medical Solutions, Erlangen, Germany). Testing began with a subjective state rating, standardized task instructions, and a blood draw. Participants were then positioned in the scanner, where two bottles—one containing 300 mL of 28% w/v glucose monohydrate (equivalent to 25% w/v glucose) and one containing water—were placed behind the head coil and connected via two plastic tubes resting on the participant’s chest (the glucose tube marked with tape).

The scan began with a 2-min baseline during which a fixation cross was shown. Participants were then instructed (via on-screen text) to grasp the marked tube, drink the glucose solution, and press a response key with their right index finger (Response Pad, Current Designs). After ingestion, they rated sweetness and palatability on visual analogue scales (0 = not at all, 100 = very). They then drank water from the second tube until the taste was neutral and confirmed completion with a key press. A fixation cross was shown for the remaining 20-min scan period. The procedure was conducted on a computer using MATLAB (R2019b) and Psychophysics Toolbox (PTB-3 v3).

The total task duration was approximately 25 min, varying slightly between participants due to self-paced drinking. Thirty-eight healthy participants were included in the analysis (19 female; mean age, 29.2 ± 6.4 years; mean fasting duration, 12.0 ± 1.7 h). The number of participants was decided using a power analysis. The power analysis estimated the required sample size to be 38, given a power of 0.85 to detect a difference between the response to glucose in the left relative to the right BLA across 20 minutes (within-subject), an alpha of 0.05, an effect size of 0.25, and a correlation among repeated measures of 0.5 (G*Power Version 3.1).

### RECORDINGS: PET imaging (humans)

This protocol was previously published^56^, and the existing dataset was reanalyzed for the present study. Ten healthy male participants underwent positron emission tomography (PET) using a Siemens HRRT scanner while receiving one of two oral stimuli: a milkshake or a tasteless noncaloric control solution. To measure dopamine release, participants were injected intravenously with 220–370 MBq [¹¹C] raclopride via a programmable syringe pump (Perfusor Compact, Braun Melsungen). Seventy percent of the tracer dose was delivered within the first minute, with the remaining 30% infused slowly over the following 59 min. Each session lasted 60 min, during which participants remained awake and motionless.

Each subject participated in two sessions (one with milkshake and one with the control solution) on separate days following a 10–11 h overnight fast.

Reanalysis of this data revealed that an experiment with this number of participants (10) would not be appropriately powered for comparison of release between right and left hemispheres. For this reason, we designed a new fMRI experiment (see above section).

### BEHAVIOR: General methods (mice)

All behavioral assays were conducted in sound-attenuated chambers (Coulbourn Habitest Modular System). Chambers were cleaned between sessions to remove residual olfactory cues. Mice were habituated for at least 1 h to the chamber—with all recording equipment and intragastric (IG) catheters attached—before the first experimental day, and for 10–20 min before each subsequent session. Both sexes were used in approximately equal numbers for all experiments. Experimenters were blinded to group identity (e.g., ChR2 vs. mCherry).

For optogenetic manipulations, light was delivered using a 473-nm DPSS laser (Shanghai Laser and Optics Century, BL473-100FC) connected via patchcords (MFP_200/220/900-0.22_2m_FCM-ZF1.25 or SBP_200/220/900-0.22_2m_FCM-GS1.0, Doric). Laser power was calibrated daily and set to 1–2 mW for inhibition and 10–15 mW for activation.

### BEHAVIOR: Flavour-nutrient conditioning (mice)

Mice underwent a 9-day flavour–nutrient conditioning paradigm (2 pre-training test days, 6 training days, 1 post-training test day) while maintained at 85–90% of initial body weight. During pre- and post-tests, mice were given 1 h two-bottle choice assays offering 0.2% saccharin solutions flavoured with 0.1% Lemon–Lime or Grape Kool-Aid (Kraft Foods). Bottle positions were randomized and counterbalanced across test days.

During training, mice were attached to IG catheters and allowed 1 h access to one flavoured solution, which triggered a 1 μL/lick IG infusion. Four counterbalanced groups associated one flavour with 8% glucose infusion and the other with water, with infusion order reversed between groups. Mouse assignment to each group was randomized, except that animals with >90% initial preference for the flavour paired with glucose infusion were excluded to avoid pre-existing bias (analyses without these mice are included in Supplementary Table 1). Mice were removed if body weight dropped below 85% or if catheters, patch cords, or cannulae disconnected during training.

Optogenetic silencing during conditioning was achieved with continuous 1–2 mW illumination in mice prepared for VTA-DA or BLA-D1R inhibition.

Optogenetic activation was achieved with 10–15 mW, 10 Hz stimulation (1 ms pulses; 2 s ON, 3 s OFF; 12 cycles) delivered after each lick following a 20 s delay, in mice prepared for VTA-DA terminal activation. In these sessions, no glucose or water was infused.

### BEHAVIOR: Conditioned flavour avoidance (mice)

Mice underwent a 3-day conditioned flavour avoidance (CFA) protocol (2 training days and 1 test day). Mice were water-deprived for 24 h before each training or testing session, with a 1-week recovery period between days. On training days, mice were given 1 h access to a 5% sucrose solution flavoured with 0.2% Orange or Cherry Kool-Aid (Kraft Foods). Prior to training, mice were naïve to both sucrose and these flavours. Ten minutes after the first lick bout, each mouse received an intraperitoneal injection of lithium chloride (100 mg/kg in saline). Mice were then returned to the chamber with continued solution access for the remainder of the session.

On the test day, mice were given simultaneous access to two bottles—one containing water and the other containing the previously flavoured sucrose solution—to assess preference. Bottle positions (front vs. back) were randomized across mice. Licks were recorded using a lickometer integrated into the Coulbourn Habitest system. Mice were excluded if patchcords or cannulae disconnected during training.

Optogenetic silencing during CFA was performed with continuous 1–2 mW illumination in mice prepared for VTA-DA or BLA-D1R inhibition.

Optogenetic activation was performed in VTA-DA terminal activation mice using 10–15 mW, 10 Hz laser pulses (1 ms pulse width; 2 s ON, 3 s OFF) delivered after the first lick following a 20 s delay, continuing for the remainder of the session. In this condition, no LiCl injection was administered.

### BEHAVIOR: Lever pressing assays (mice)

Mice were trained to self-stimulate VTA-DA terminals using optogenetic activation. Stimulation was delivered through implanted cannulae connected to a 10–15 mW, 473-nm laser as described above. Mice underwent a 3-day lever-pressing protocol in which they had access to two levers for 1 h per day. Pressing one lever triggered terminal stimulation (20 Hz, 1 ms pulse width, 5 s total). The position of the active lever (front vs. back) was randomized on the first day and held constant thereafter. Mice were excluded if patchcords or cannulae disconnected during any session.

### BEHAVIOR: Feeding assays (mice)

In a set of related assays, mice were given access to flavoured Ensure solutions during optogenetic inhibition or activation. On four separate days, mice were provided 30 min access to a flavoured Ensure solution following 1 h of water and food deprivation. Two sessions were conducted with the laser off and two with the laser on, in counterbalanced order. Consumption was averaged across replicate “laser on” and “laser off” sessions. Testing time was kept consistent (±30 min) to control for circadian effects.

Optogenetic silencing during feeding was achieved with continuous 1–2 mW illumination in mice prepared for VTA-DA or BLA-D1R inhibition. The laser remained on throughout access to a vanilla Ensure solution. If patchcords or cannulae disconnected during a “laser on” session, data from that day were discarded and the session repeated if possible.

Optogenetic activation during feeding was performed in mice prepared for VTA-DA terminal activation. Three flavoured Ensure solutions (vanilla, chocolate, strawberry) were tested in separate assays:

- Continuous stimulation – 10 Hz, 1 ms pulses (2 s ON, 3 s OFF) delivered during access to vanilla Ensure.
- Lick-terminated stimulation – same parameters, but laser turned off for 2 s following each lick (chocolate Ensure).
- Lick-triggered stimulation – same parameters, but laser turned on for 2 s after each lick (strawberry Ensure or water).

If cannulae disconnected during any “laser on” day, data were discarded and the session re-run if possible.

### ANALYSIS: Recordings (mice)

All analyses were performed using custom MATLAB scripts (R2023a). Unless otherwise stated, data are presented as mean ± SEM, with shaded areas in line plots indicating SEM or 95% confidence intervals for regression fits. Except where noted (e.g., linear regressions, Kolmogorov–Smirnov tests), all statistical comparisons used nonparametric two-sided permutation tests (100,000 iterations or exact when possible). One-sided tests were applied sparingly, in cases where directionality was pre-specified, specifically in two cases: (1) in conditioned flavour avoidance experiments, where activation or LiCl injection could only decrease sucrose preference since it can be near 100% at baseline (2) in self-stimulation experiments, where activation could only increase lever pressing since it can be near 0 at baseline.

### ANALYSIS: Time windows (mice)

Activity was analyzed across three standardized time windows:

#### Oral phase

0–5 s after the first lick in a bout. Lick bouts were defined as clusters of licks separated by ≤2 s and lasting ≥2 s. Only the first bout was analyzed (except in 2-photon recordings), as post-ingestive signals are minimal during this period. Given that mice consume ∼9.6 µL/s^64^ and swallow at ∼1 Hz^65^, a 5 s bout maximally represents ∼38 µL swallowed (≈10% of baseline stomach volume^66^), consistent with minimal GI feedback.

#### IG phase

20–600 s after the start of intragastric (IG) infusion, corresponding to the arrival of solution in the stomach (∼20 s transit) and continuing through the 10 min infusion period.

#### Post-IG phase

1200–3000 s after infusion onset (10–40 min post-infusion), allowing for fluid clearance from the stomach and intestinal transit (which based on oral gavage studies exceeds 5 min^67^).

#### Hormone phase

0–300 s after intraperitoneal (IP) injection. Useful for quantifying immediate responses to systemic injection.

#### Malaise phase

300–1800 s after IP injection. This time window was only used for LPS responses, owing to delayed action.

### ANALYSIS: Catheter transit time (mice)

To determine catheter transit time, we measured the time it took for Ensure and water to pass through catheters of the same lengths as those implanted in mice used for recordings. Transit times for water were consistent with values previously reported by our lab^68^. Transit times for Ensure were slightly longer, likely reflecting its higher viscosity.

### ANALYSIS: Photometry (mice)

Fluorescence signals were acquired at isosbestic (420 nm) and activity-dependent (470 nm) wavelengths. The activity-dependent signal was normalized to the isosbestic trace to remove motion artefacts using ΔF/F₀ = (F − F₀)/F₀, where F₀ was predicted from a linear regression of the two signals during the 10 min baseline. For IG and IP manipulations, each recording was further normalized by subtracting the corresponding sham or saline trace from the same mouse. Un-subtracted traces are shown in example figures, while subtracted traces were used for statistics and summary plots.

To account for relative differences in DA axon density, we normalized photometry traces by z-scoring relative to baseline activity during drinking in Fig. 2. The raw ΔF/F₀ activity for this experiment can be found in the supplement (Extended Fig. 3).

### ANALYSIS: Single-cell imaging (mice)

Neural activity traces were extracted after CNMF-E segmentation (see Recording Methods) and z-scored relative to baseline activity. Lick timing was aligned to neural traces via TTL signals. For miniscope data, only the first lick bout was analyzed; for 2-photon recordings, all bouts were combined (typically <5 s each, separated by >20 s).

#### Sham sessions and control distributions

To control for baseline drift and intrinsic excitability differences across neurons and animals, we recorded “sham” sessions in which mice were placed in the behavioral chamber but received no stimulation, infusion, or reward. These sessions were used to generate empirical distributions of spontaneous neuronal activity for each time window. For lick-related analyses, TTL timestamps from real licking sessions (e.g., Ensure or sucrose) were applied to sham recordings to create matched sham lick responses.

#### Activated and inhibited thresholds

Neurons were classified as activated or inhibited using empirical thresholds derived from sham distributions, rather than fixed z-score cutoffs. A neuron was considered activated if its z-scored activity exceeded the 95th percentile of its sham distribution and inhibited if below the 5th percentile (α=0.05).

This empirical approach was adopted because fixed ±1z thresholds, commonly used in the field (and in our prior work^25^), assume homogeneous baseline variance across neuron types and timescales—an assumption that does not hold for mixed populations or long recordings. By contrast, this sham-based method defines activation relative to each neuron population’s own spontaneous activity, thereby controlling for intrinsic excitability, drift, and noise.

Because baseline variance differed across time windows, independent thresholds were computed for each phase (e.g., IG vs. post-IG). For instance, 5% of VTA-DA/CCK neurons exceeded +0.54 z during sham IG infusions and +0.69 z during sham post-IG periods. Distinct z-score thresholds were similarly derived for each neuron group (e.g., VTA-DA/LepR vs. VTA-DA/CCK) to control for differences in baseline excitability. Sham-derived distributions for licking responses were used in the same way to determine activation and inhibition cutoffs.

Cross-registration for neurons across days and conditions was performed manually. This was achievable due to the sparsity of VTA-DA/CCK, VTA-DA/LepR, and aBLA-D1R neuron subpopulations.

Rise times for dopamine release were calculated from individual traces during IG infusions using a window from 1 minute before infusion onset to 5 minutes after. For lick response rise times, the window was the 5 seconds before and after lick bout onset. Traces were fitted with a sigmoid curve, and the time at which the fit reached 5% of its final value was defined as T5%. The first time point exceeding baseline fluorescence by 3 standard deviations (σ, defined during baseline) was defined as T3σ. Both metrics are shown unadjusted in Supplementary Table 2.

For comparisons reported in the text, catheter transit time was subtracted to yield adjusted rise times. Adjusted T5% and T_3σ_ values were averaged to compute the reported mean rise time, with standard errors combined as 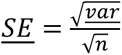. Individual neuron recordings could not always use both rise-time measures: IG responses did not consistently follow a sigmoid function (so only T3σ was used), whereas lick-related responses rarely exceeded 3σ (so only T5% was used). Consequently, rise times for neural activity and dopamine release were compared using the same metric within each analysis (e.g., IG responses using T3σ).

Fall times were calculated analogously, using a window from 5 minutes before the end of infusion to 40 minutes after infusion offset.

Peak times for dopamine release during licking was determined from the first lick bout (0–30 s window from bout onset), defined as the time of maximal fluorescence within this period. For aBLA neuron activity, peak timing was measured in a shorter 0–5 s window after lick-bout onset, corresponding to the briefer access period.

Statistical corrections for between-subject variance were utilized when noted, with a linear mixed-effects model that treated individual mice as random effects. Intra-class correlations (ICC) were computed as 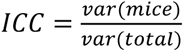 to quantify variance attributable to inter-animal differences. Where appropriate (e.g., Extended Fig. 8l), activity traces were also averaged by mouse to visualize ICC effects.

Clustering analyses were performed using principal component analysis (PCA) and k-means clustering. Traces were first downsampled (to 1 Hz for IG responses or 4 Hz for lick responses) and normalized by their absolute maximum. PCA was performed using MATLAB’s built-in pca function with centering by singular value decomposition. The number of components plotted was determined by the elbow criterion on the scree plot (when unclear, the first 2 PCs were plotted alone). k-means clustering was implemented using MATLAB’s kmeans function, with cluster number chosen by the silhouette method. Cluster assignments were then visualized using the unnormalized data.

### ANALYSIS: fMRI recordings (humans)

All fMRI preprocessing and analyses were performed using SPM12 (Wellcome Trust Centre for Neuroimaging) and FMRIB software library (FSL, version 5.0.10) implemented in MATLAB (R2022b). Functional data were motion- and distortion-corrected (mcflirt^69^, topup^70^), co-registered to individual structural images, normalized to MNI space using unified segmentation, and spatially smoothed with a 6 mm FWHM Gaussian kernel. The first 10 volumes were discarded to allow for magnetic stabilization.

At the first level of analysis, we modeled the Drink Task using a mass-univariate general linear model (GLM). The baseline (2 min) and ten consecutive 2 min post-ingestion time bins were modeled as separate boxcar regressors convolved with the canonical hemodynamic response function and its time and dispersion derivatives. The drinking and rinsing phases were modeled as an additional regressor of variable duration. Nuisance regressors included 24 motion parameters (current and preceding volume and their squares^70^), mean ventricular CSF signal, and motion-outlier volumes (> 1 mm framewise displacement^71^). Low-frequency drifts were high-pass filtered at 128 s. Ten contrasts (post-ingestion > baseline) were computed for each subject.

Second-level analysis used a flexible factorial GLM including subject and condition factors, with sex, hunger rating, outside temperature, and insulin sensitivity (HOMA-IR) as covariates. We identified voxels that were activated in response to glucose relative to baseline. Significance was assessed within anatomically defined left and right BLA masks^72^ (threshold = 0.1) at p < 0.05, FWE-corrected at the peak level (initial voxel threshold = 0.05 uncorrected). For comparing activity across hemispheres, we fitted a mixed-effect model using BLA side (L vs. R) and time as a fixed effects, with a random intercept for the individual subject. Linear mixed effects models and type III analyses of variance were fitted via the restricted maximum likelihood and Satterthwaite’s methods, respectively, using the lmerTest package (version 3.1-3) in RStudio (2023.06.0+421, 2009-2023 Posit Software, PBC).

We aimed to directly test whether there is a lateralized response in the BLA to glucose consumption. To this end, we first extracted single-subject values from the left and right BLA peaks that were maximally responsive to glucose consumption. We then fitted a linear mixed-effects model with neural response to glucose as the dependent variable, and BLA location (right vs left), time bin (1:10-2 min), and their interaction as fixed effects, with a random intercept for subject ID.

### ANALYSIS: Behavior (mice)

Behavioral data were analyzed from Coulbourn system outputs of licking and lever pressing. Lick data are shown as cumulative totals or as flavour preferences, calculated as the percent preference for the flavour paired with neuronal stimulation, glucose infusion, or LiCl injection: A/(A+B), where A is the total number of licks for one flavour and B for the other.

Lever-pressing behavior is shown as the difference between active and inactive lever presses. Statistical tests were corrected for multiple comparisons using the Benjamini–Hochberg procedure, where n denotes the number of brain regions tested (see Fig. 4 and Extended Data Fig. 7 and Supplementary Table 2).

### ANALYSIS: Histology (mice only)

Mice were transcardially perfused with phosphate-buffered saline (PBS) followed by 10% formalin. Brains were post-fixed overnight at 4 °C, cryoprotected in 15% and then 30% sucrose (24 h each), embedded in OCT (Tissue-Tek), and sectioned on a cryostat (50 μm).

For immunostaining, free-floating sections were washed (3 × 5 min in PBS-T, 0.1% Triton X-100), blocked for 2 h at room temperature (2% normal goat serum + 3% BSA in PBS-T), and incubated for 12–48 h at 4 °C with primary antibodies in blocking solution. Sections were washed, incubated with fluorescent secondary antibodies for 2 h at room temperature, washed again, and mounted with DAPI Fluoromount-G (Southern Biotech). Images were acquired on an Eclipse Ti2 inverted microscope (Nikon) and minimally processed in ImageJ (v1.53t).

Primary and secondary antibodies included: chicken anti-GFP (Abcam ab13970, 1:500) + goat anti-chicken Alexa 488 (Abcam ab150169, 1:500), and rabbit anti-RFP (Abcam ab124754, 1:500) + goat anti-rabbit Alexa 568 (Abcam ab175471, 1:500).

Fiber placement and viral spread were verified visually to generate anatomical placement maps; median A–P slices are shown for each group. Because ML/DV coordinates vary along the A–P axis, some tracts on this median slice may appear slightly displaced. Mice with clear off-target fiber placements were excluded from analysis.

Pseudorabies retrograde tracing. Labeling from pseudorabies retrograde tracing was quantified using the QUINT semi-automated whole-brain analysis pipeline^73^, which integrates image registration, segmentation, and region-based cell counting across the Allen CCFv3 atlas. This produced quantitative “load” measurements (GFP⁺ cells per unit area) for each of the 1,328 annotated brain regions. Segmentation and registration accuracy were manually validated in representative sections from each animal. Regions smaller than 100 pixels in any dimension and non-gray-matter annotations were excluded to avoid artefactual load estimates. Loads were normalized to VTA labeling to control for viral transfection efficiency, ranked by difference following left vs right BLA injections, and visualized as force-directed graphs, dendrograms, and heat maps using connectivity matrices (0–5 scale) derived from literature curation.

Rabies retrograde tracing. GFP⁺ cells were manually counted from five representative midbrain sections in each of three mice.

### ANALYSIS: Graphing

For visualization, fluorescence and activity traces were down-sampled and smoothed using a moving-average filter (kernel size = 2). Long-duration traces (e.g., IG responses) were down-sampled to 0.1–0.2 Hz, whereas short-duration traces (e.g., licking responses) were down-sampled to 1 Hz. Principal component (PC) space plots of licking activity were maintained at 4 Hz to preserve temporal resolution.

In some cases, the same dataset was visualized in multiple figures for distinct comparisons (e.g., dopamine responses during Ensure IG infusion in left BLA are shown in Fig. 2a, and Extended Figs. 2g, 10a to compare responses with striatum, water infusion, and contralateral BLA, respectively).

### Statistics and Reproducibility

We confirmed our main findings by repeating recordings and behavioral experiments in multiple animals, and repeating all recordings in the same animal whenever possible. All attempts at replication for experiments shown in the manuscript were successful. In behavioral experiments (except for learning assays) as well as photometry recordings in the manuscript, two replicates per mouse per condition were used. The data reports an average of these replicates. In photometry recordings, two replicates were performed per mouse per condition. The data reports an average of these replicates as well. In other single-cell recordings, replication of the main findings were performed for the main findings, but only one replicate per animal is shown in the manuscript as averaging the activity of single neurons removes important dynamics like spiking.

## Data Availability

The data used to generate the main findings of this study have been publicly deposited at Dryad at https://doi.org/10.5061/dryad.h9w0vt4x2.

## Code Availability

No custom software or original algorithms were developed for use in this study. Any of the findings can be replicated using established software and the deposited data.

## Acknowledgements

We thank K. Yackle and I. Bachmutsky for providing AAV-Con/Fon-ChR2 viruses. We thank A. MacDonald for his help establishing the vagotomy method.

## Contributions

J.C.R.G. and Z.A.K. conceptualized the study and wrote the manuscript with input from D.M.S. and M.T. All experiments done in mice were designed, analyzed, and interpreted by J.C.R.G. and Z.A.K. Intragastric surgeries were performed by J.C.R.G. and L.Q.; all other surgeries were performed by J.C.R.G. Mouse behavior and in vivo imaging studies were performed by J.C.R.G. and Q.L. Lateralized network tracing studies were performed by J.C.R.G., A.M.H., J.Z., and Z.L. All other histological experiments were performed by J.C.R.G., A.M.H. J.C., and V.U. All experiments done in humans were designed, run, analyzed, and interpreted by H.B., B.K., D.M.S., and M.T.

## Additional Information

Correspondence should be addressed to Zachary A. Knight at.

## Extended Figure Legends

**Extended Data Fig. 1.**
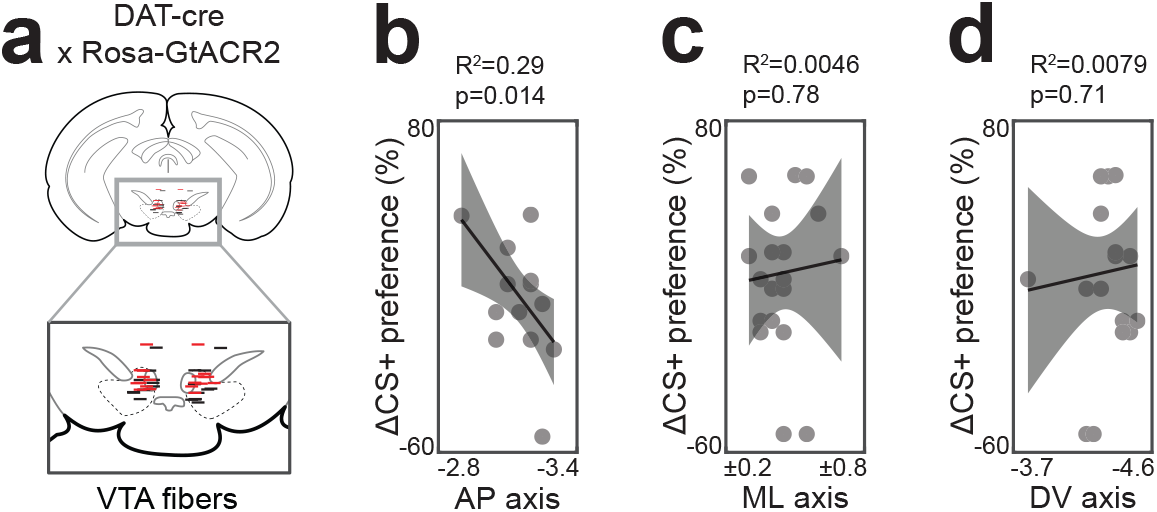
Fiber localization for VTA-DA inhibition experiments. a, Fiber placement in VTA, shown with median slice along anteroposterior (AP) axis (black=littermate controls, red=GtACR). b, Mice with fibers above posterior VTA showed greater inhibition of flavour learning, consistent with the location of the BLA projecting population (Lammel et al.,2008). c, Fiber placement along the mediolateral (ML) axis did not affect learning, as deviations would cancel within animals due to the fixed bilateral spacing of implants. d, Fiber placement along the dorsoventral (DV) axis did not affect learning. ns P > 0.05; *P < 0.05; **P < 0.01; ***P < 0.001. Shaded areas show standard error of the mean (SEM). See Supplementary Table 1 for statistical tests, n, and p-values.

**Extended Data Figure 2.**
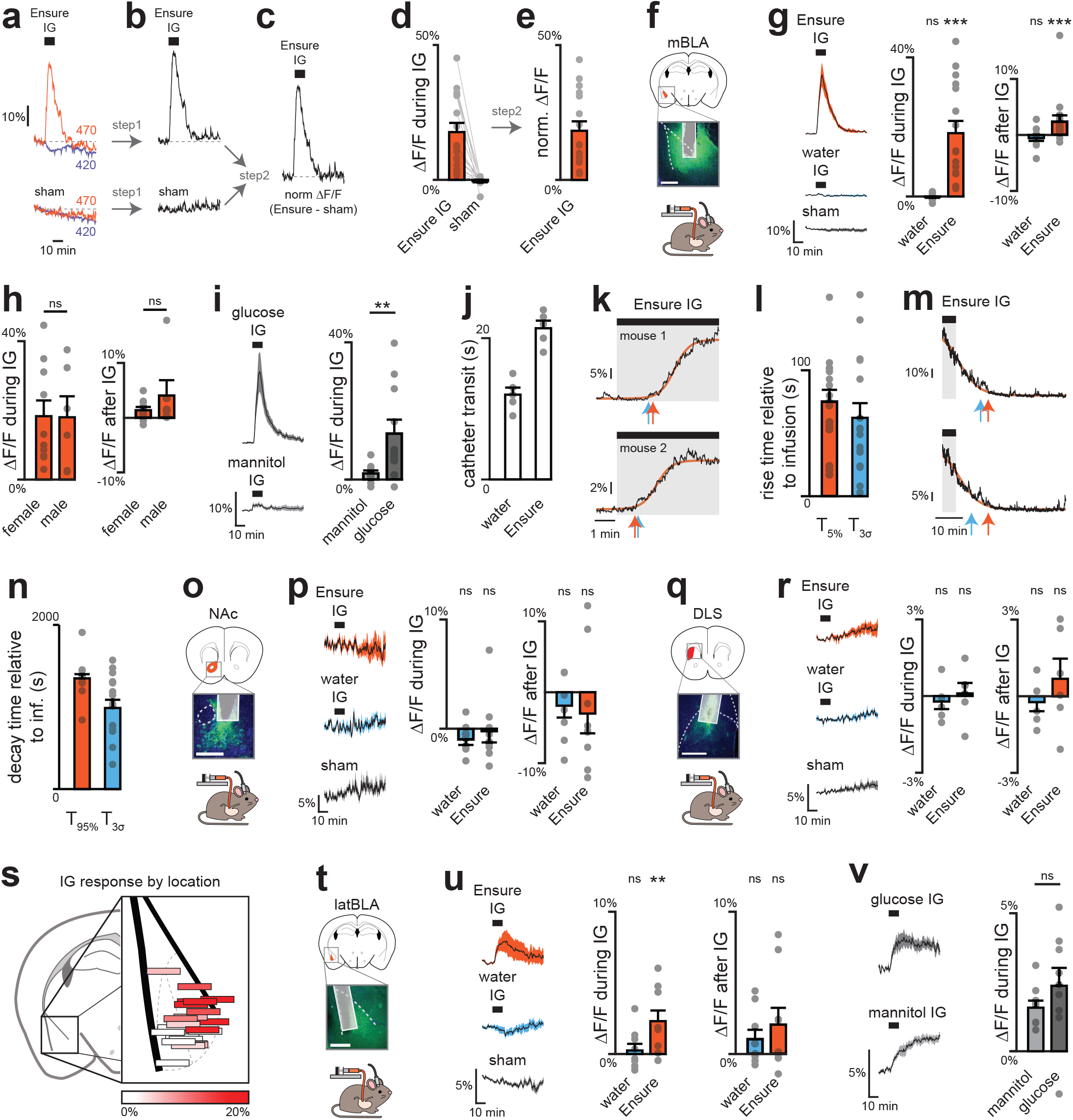
Dopamine dynamics across brain regions during IG infusions. a–e, Two-step normalization procedure. Step 1: fluorescence traces normalized to activity-independent signal (420 nm), removing movement artefacts. Step 2: traces during IG or IP manipulations further normalized to sham infusion or saline injection, correcting for bleaching and long-term drift. Outside this figure, only step 1 normalization was applied to activity traces, and step 2 was applied only to summary plots. a, Example raw fluorescence traces from a single mBLA GRAB-DA site during Ensure IG (top) or sham (bottom). b, Dopamine-dependent signal after 470 nm traces were normalized to 420 nm. c, Further normalization by subtracting sham traces. d, Summary of mBLA dopamine release during Ensure IG vs sham after step 1. e, Summary of the same data after both steps. f, GRAB-DA recordings in mBLA; dotted lines mark external capsule (scale bar, 0.5 mm). g, Mean traces (left) and summary plots (middle, right) showing greater mBLA DA release during and after Ensure infusion. h, No sex differences in mBLA DA release during (left) or after (right) Ensure infusion. i, Glucose, but not mannitol, increases mBLA DA release. j, Transit times for IG infusions: water 12 ± 1 s; Ensure 21 ± 1 s. k, Example traces with sigmoid fits to estimate T5% (orange) and T3σ (blue). l, Summary rise times relative to infusion onset, corrected for transit time (from e): T5% = 54 ± 9 s; T3σ = 41 ± 12 s; mean = 48 ± 11 s. m, Example traces with sigmoid fits to estimate T95% (orange) and T3σ (blue). n, Summary decay times relative to infusion end: T95% = 749 ± 51 s; T3σ = 388 ± 100 s; mean = 569 ± 79 s. o, GRAB-DA recordings in NAc core; dotted lines mark anterior commissure (scale bar, 0.5 mm). p, No change in NAc DA release with Ensure or water. q, GRAB-DA recordings in DLS; dotted lines mark corpus callosum (scale bar, 0.5 mm). r, No change in DLS DA release with Ensure or water. s, Reconstructed fiber placements in BLA (color scale = ΔF/F during Ensure IG), showing greater DA release in magnocellular vs parvocellular BLA. t, GRAB-DA recordings in lateral BLA; dotted lines mark external capsule (scale bar, 0.5 mm). u, Lateral BLA shows increased DA release during Ensure infusion, but not after. v, Both glucose and mannitol increase DA release in lateral BLA. ns P > 0.05; *P < 0.05; **P < 0.01; ***P < 0.001. Shaded areas in mean traces and error bars in bar plots show SEM. See Supplementary Table 2 for statistical tests, n, and p-values.

**Extended Data Figure 3.**
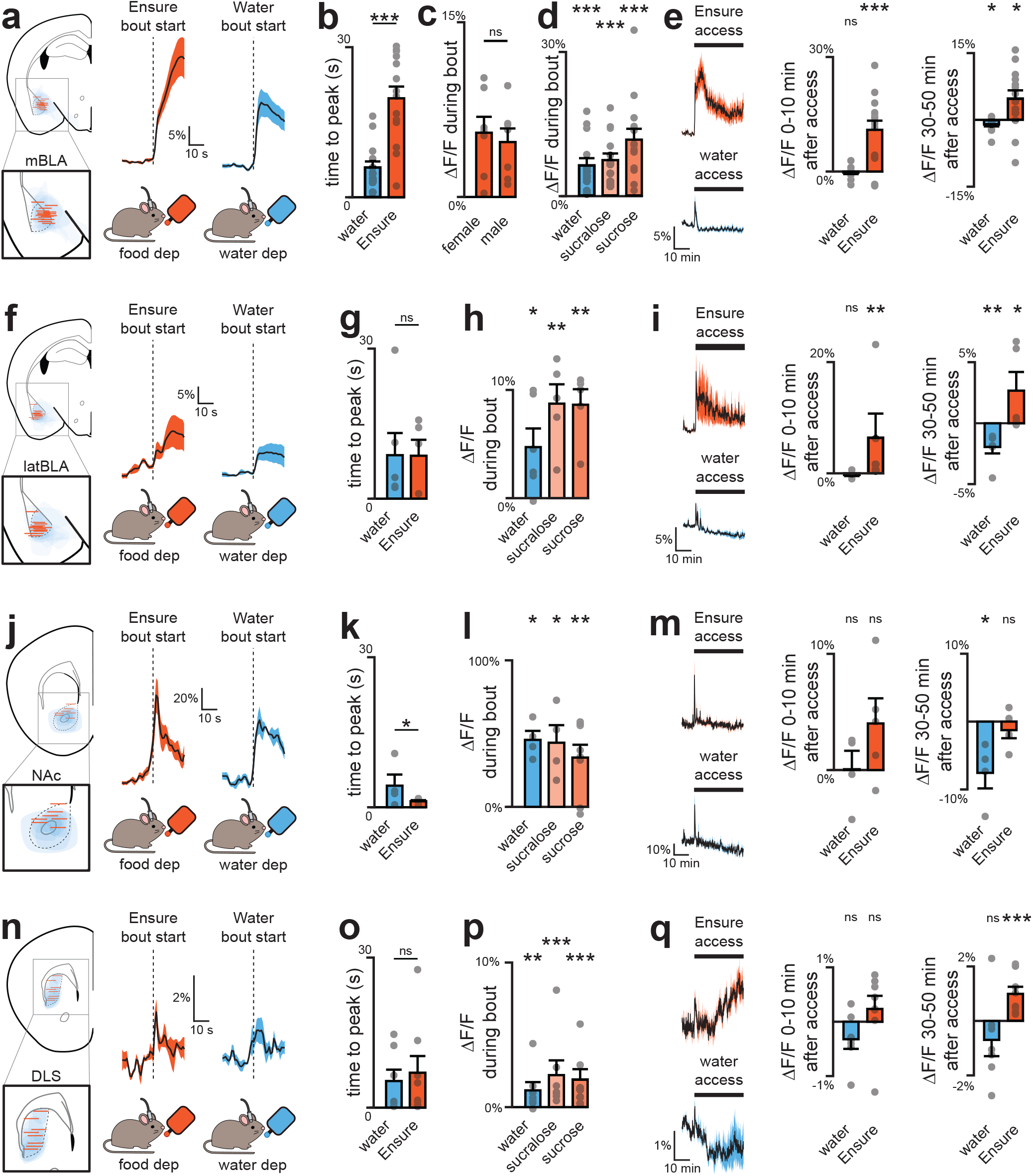
Dopamine dynamics across brain regions during natural consumption. a, Fiber placement and viral spread for mBLA recordings (left). Mean GRAB-DA responses during first lick bout of Ensure and water (middle, right). b, mBLA DA peak time is delayed following Ensure vs water. c, No sex differences in mBLA DA release during first Ensure lick bout. d, Summary plots of mBLA DA release during first lick bouts. e, Mean traces (left) and summary plots (middle, right) showing long-term mBLA DA increases during Ensure consumption. f, Fiber placement and viral spread for lateral BLA recordings (left). Mean GRAB-DA responses during first lick bout of Ensure and water (middle, right). g, Peak time of lateral BLA DA release. h, Summary plots of lateral BLA DA release during first lick bouts. i, Mean traces (left) and summary plots (middle, right) showing long-term lateral BLA DA increases during Ensure consumption. j, Fiber placement and viral spread for NAc recordings (left). Mean GRAB-DA responses during first lick bout of Ensure and water (middle, right). k, Peak time of NAc DA release. l, Summary plots of NAc DA release during first lick bouts. m, Mean traces (left) and summary plots (middle, right) showing no long-term NAc DA increase during Ensure consumption. n, Fiber placement and viral spread for DLS recordings (left). Mean GRAB-DA responses during first lick bout of Ensure and water (middle, right). o, Peak time of DLS DA release. p, Summary plots of DLS DA release during first lick bouts. q, Traces (left) and summary plots (middle, right) showing long-term DLS DA increases during Ensure consumption. ns P > 0.05; *P < 0.05; **P < 0.01; ***P < 0.001. Shaded areas in mean traces and error bars in bar plots show SEM. See Supplementary Table 2 for statistical tests, n, and p-values.

**Extended Data Fig. 4.**
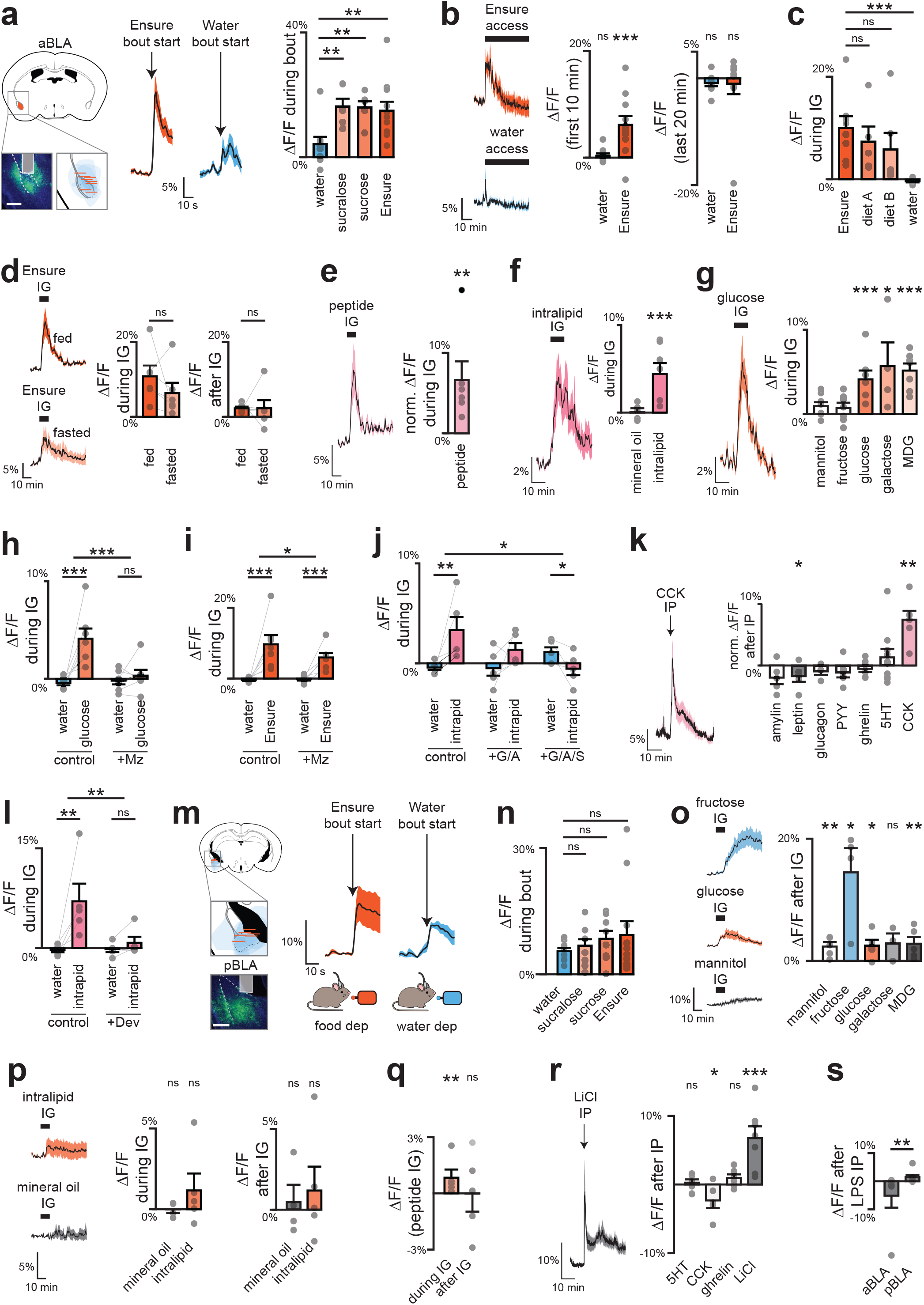
| Dopamine release depends on nutrients in anterior, but not posterior BLA. a–l, Dopamine (DA) recordings in posterior BLA (pBLA). a, Fiber placement and viral spread for aBLA recordings (left; scale bar, 0.5 mm). DA dynamics during first lick bout of water and Ensure (middle) with summary plot showing greater release to sweet solutions vs water (right). b, Traces (left) and summary plots (right) of DA release during Ensure or water access. c, Summary plots during access to calorie-matched diets: diet A (55%/14%/31% sugar/fat/protein, matches Ensure) and diet B (20%/20%/60%, matches mouse preference). d, Traces (left) and summary plots (right) showing no difference in DA response to Ensure IG in fed vs fasted mice. e, Traces (left) and summary plots (right) of DA release during peptide IG. f, Traces (left) and summary plots (right) of DA release during intralipid vs mineral oil IG. g, Traces (left) and summary plots (right) of DA responses to glucose and sugar analogs during IG. h, Summary plot of DA responses during water or glucose IG with or without SGLT1 antagonist mizagliflozin (Mz). i, Summary plot of DA responses during water or Ensure IG with or without mizagliflozin. j, Summary plot of DA responses during water or intralipid IG with or without antagonists of GPR40 (GW1100, G), GPR120 (AH7614, A), or CD36 (sulfosuc-cinimidyl oleate, S). k, Traces (left) and summary plots (right) of DA release during IP injection of CCK and other hormones. l, Summary plot of DA release during water or intralipid IG with or without prior CCK antagonist devazepide (Dev). m–s, Dopamine (DA) recordings in posterior BLA (pBLA). m, (Left) Fiber placement and viral spread for pBLA recordings (scale bar, 0.5 mm). (Right) DA dynamics during first lick bout of water and Ensure. n, No difference in DA release during bouts of water vs nutrients. o, Traces (left) and summary plots (right) showing increased DA release after IG sugar infusion, independent of SGLT1. p, Traces (left) and summary plots (right) showing no DA increase after nutritive or non-nutritive fat infusions. q, Small but significant DA release during peptide infusion. r, Traces (left) and summary plots (right) showing no DA response to feeding hormones, but significant release to aversive LiCl (IP). s, Increased pBLA DA and decreased aBLA DA following aversive LPS injection. ns P > 0.05; *P < 0.05; **P < 0.01; ***P < 0.001. Shaded areas in mean traces and error bars in bar plots show SEM. See Supplementary Table 2 for statistical tests, n, and p-values.

**Extended Figure 5.**
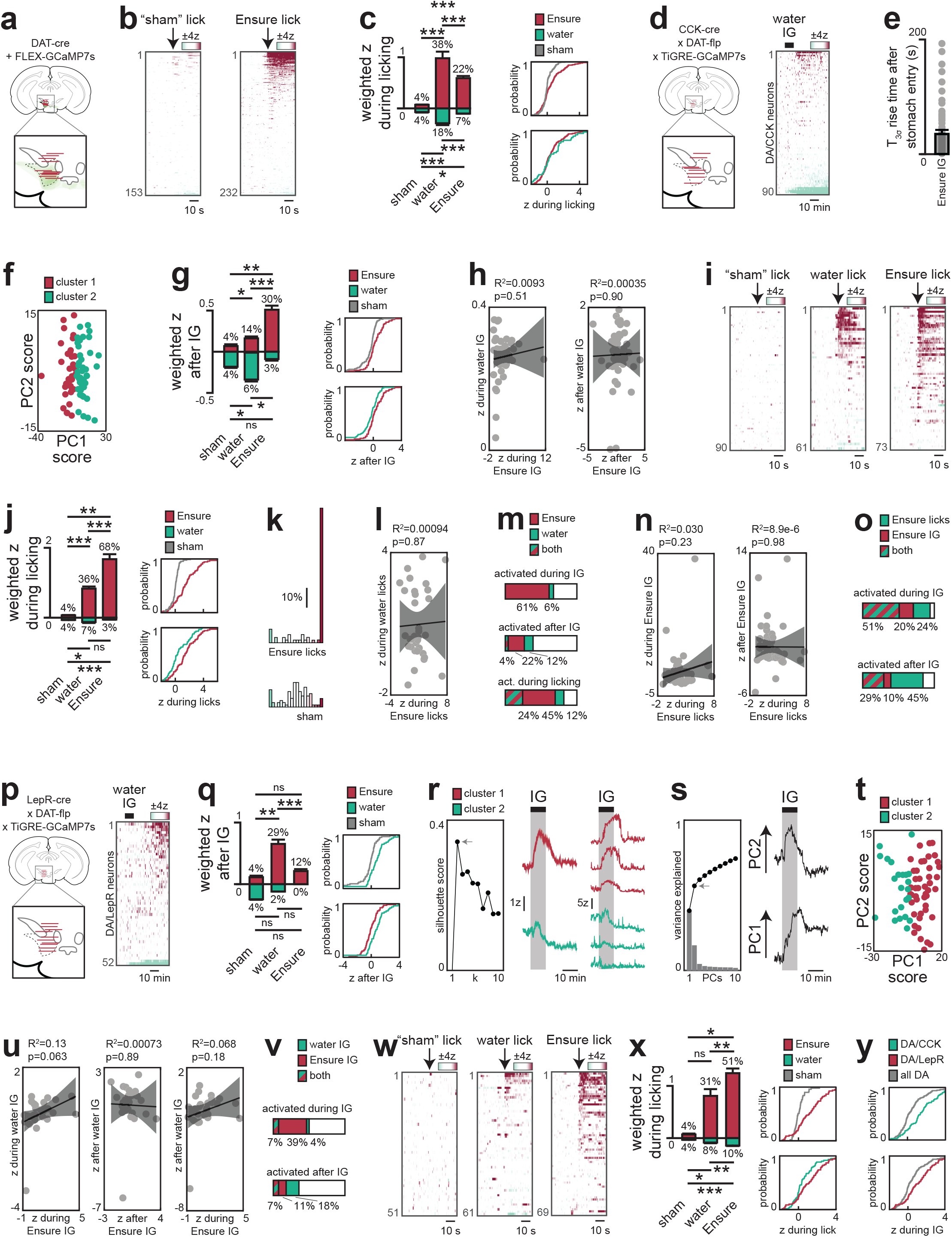
DA subpopulation responses across different stages of ingestion. a–c, Single-cell imaging of all VTA-DA neurons. a, Fiber placement and viral spread for imaging all VTA-DA neurons. b, Dynamics during consumption of Ensure or sham (see Methods). c, Population-weighted activity of activated (magenta) and inhibited (mint) neurons (left) and cumulative distributions (right) showing increased activity during the first Ensure bout (vs sham, p = 2.6e-6; vs water, p = 0.24). d–o, Imaging of VTA-DA/CCK neurons. d, Fiber placement for imaging VTA-DA/CCK neurons (left). Dynamics following water infusion (right). e, Activity rose 36±7 s after Ensure reached the stomach. f, PC1 separates transient vs prolonged subtypes from k-means clustering. g, Population-weighted activity (left) and cumulative distributions (right) showing increased activity after Ensure infusion (vs sham, p = 1.7e-4; vs water, p = 0.038). h, Correlation analyses indicate distinct responses to water vs Ensure infusion. i, Dynamics during consumption of Ensure, water or sham. j, Population-weighted activity (left) and cumulative distributions (right) confirm greater responses during Ensure bouts (vs sham, p = 2.0e-15; vs water, p = 0.0013). k, Binned activity during Ensure bouts. l, Correlation analyses indicate distinct responses to water vs Ensure bouts. m, Overlap of neurons responding to water and Ensure. n, Correlation analyses indicate distinct responses during IG vs natural consumption. o, Overlap of neurons responding during IG and natural consumption. p–z, Imaging of VTA-DA/LepR neurons. p, (Left) Fiber placement for recording VTA-DA/LepR neurons. (Right) Dynamics during water infusion. q, Population-weighted activity (left) and cumulative distributions (right) show activation after water but not Ensure infusion (water vs sham, p = 6.9e-5; water vs Ensure, p = 0.0042; Ensure vs sham, p = 0.33). r, K-means clustering reveals prolonged (purple) and transient (green) subpopulations, shown with mean and representative single-neuron activity. s, Principal component analysis; the first two PCs explain 56% of variance and correspond to activity changes during and after infusion. t, PC1 separates transient vs prolonged subtypes from k-means clustering. u, Correlation analyses indicate distinct responses to water vs Ensure infusion. v, Overlap of neurons responding to water and Ensure infusions. w, Dynamics during consumption of Ensure, water or sham. x, Population-weighted activity (left) and cumulative distributions (right) show increased activity during Ensure bouts (vs sham, 1.2e-6; vs water, p=0.045). y, Cumulative distributions across subpopulations showed robust activation during Ensure consumption in both DA/CCK (p = 4.2e-11) and DA/LepR (p = 2.3e-12) groups. ns > 0.05; *P < 0.05; **P < 0.01; ***P < 0.001; ****P < 0.0001. Shaded areas show SEM in mean traces and correlation plots. Error bars show SEM in bar plots. See Supplementary Table 3 for statistical tests, n, and p-values.

**Extended Figure 6.**
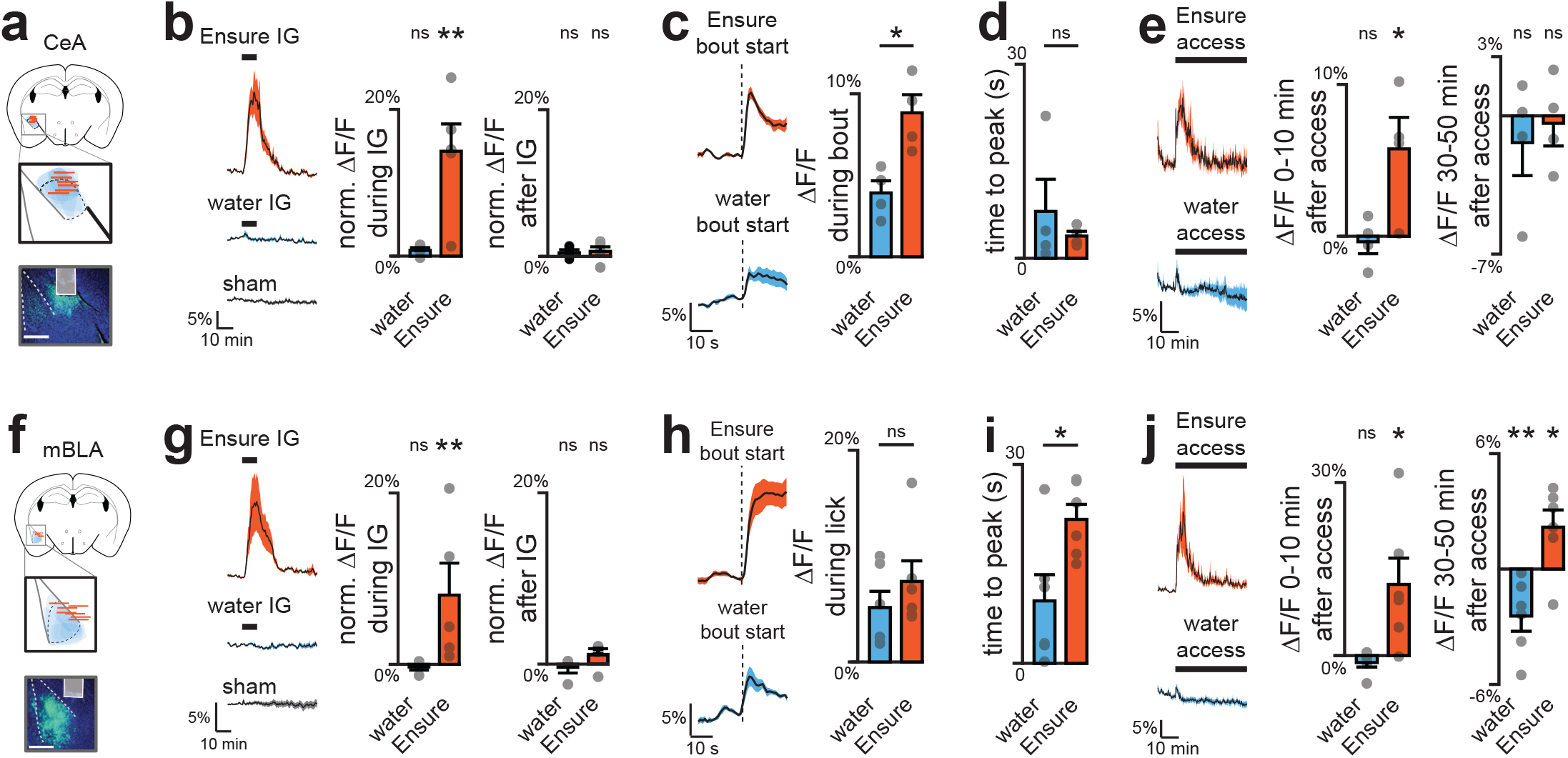
DA release in neighboring mBLA and CeA during ingestion. a–e, Fiber placement and viral spread for recording DA release onto CeA-VGAT neurons (a).DA increased during Ensure IG infusion compared to controls (b) and during the first lick bout (c), with no difference in peak timing across solutions (d). Mean activity remained elevated throughout consumption (e). f–j, Fiber placement and viral spread for recording DA release onto mBLA-Vglut neurons (f). DA increased during Ensure IG infusion (g) and during the first lick bout (h), with peak release delayed for Ensure vs water (i). Mean activity remained elevated throughout consumption (j). ns > 0.05; *P < 0.05; **P < 0.01; ***P < 0.001; ****P < 0.0001. Shaded areas in mean traces and error bars in bar plots show SEM. See Supplementary Table 2 for statistical tests, n, and p-values.

**Extended Data Figure 7.**
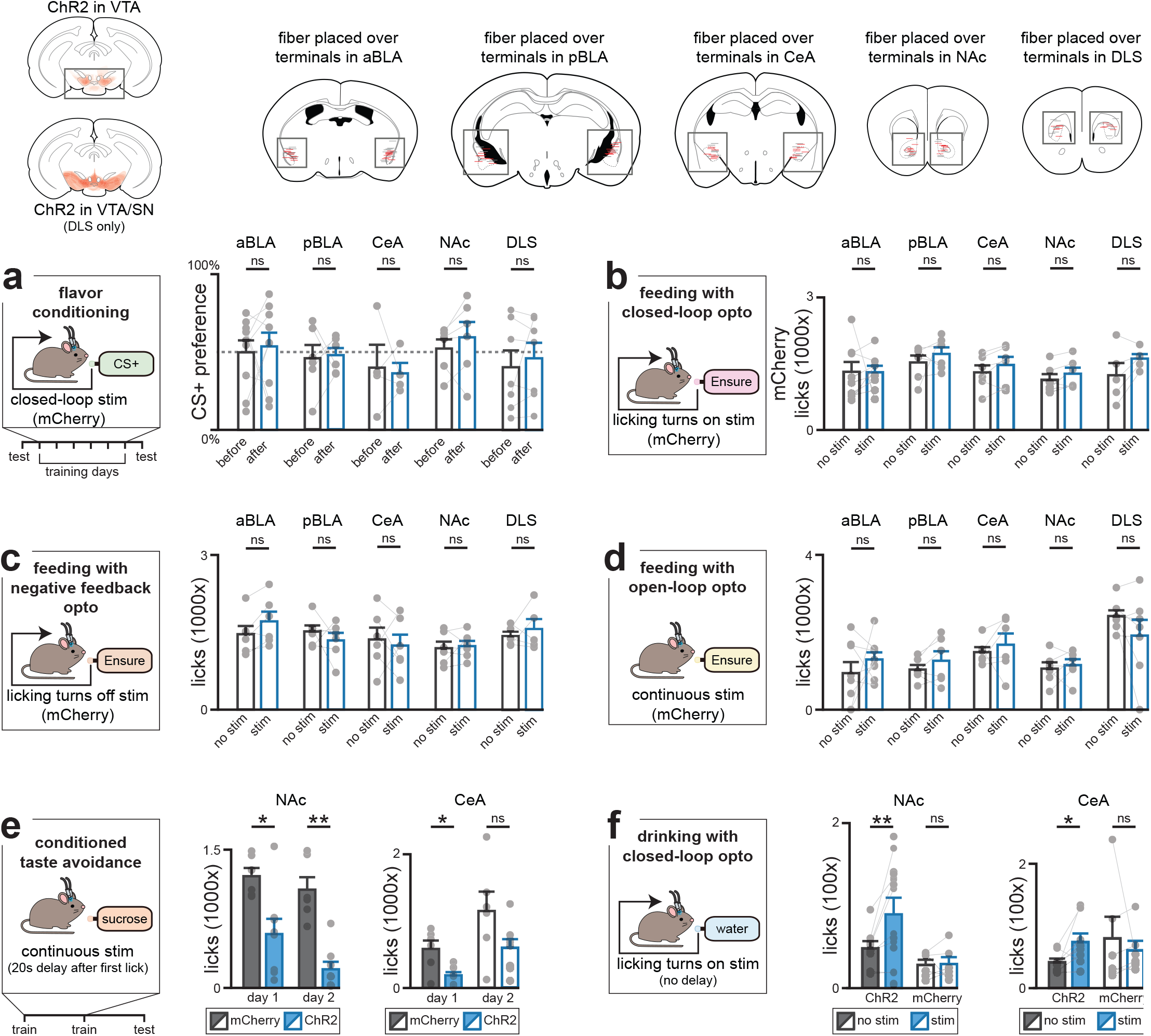
Controls for dopamine projection experiments. (Top) Viral expression of ChR2 in VTA-DA neurons (or all midbrain DA neurons for DLS terminal experiments) and fiber placement above projection targets (grey=mCherry, red=ChR2; see Fig. 4 for insets). a, Flavour conditioning assay (mCherry controls; Fig. 4a). b, Lick-activated stimulation (mCherry controls; Fig. 4d). c, Lick-terminated stimulation (mCherry controls; Fig. 4e). d, Continuous stimulation (mCherry controls; Fig. 4f). e, Sucrose consumption during CTA training. f, Lick-activated stimulation with water substituted for Ensure. *P < 0.05; **P < 0.01; ***P < 0.001, after Benjamini–Hochberg correction. Error bars show SEM. See Supplementary Table 1 for statistical tests, n, and p-values.

**Extended Figure 8.**
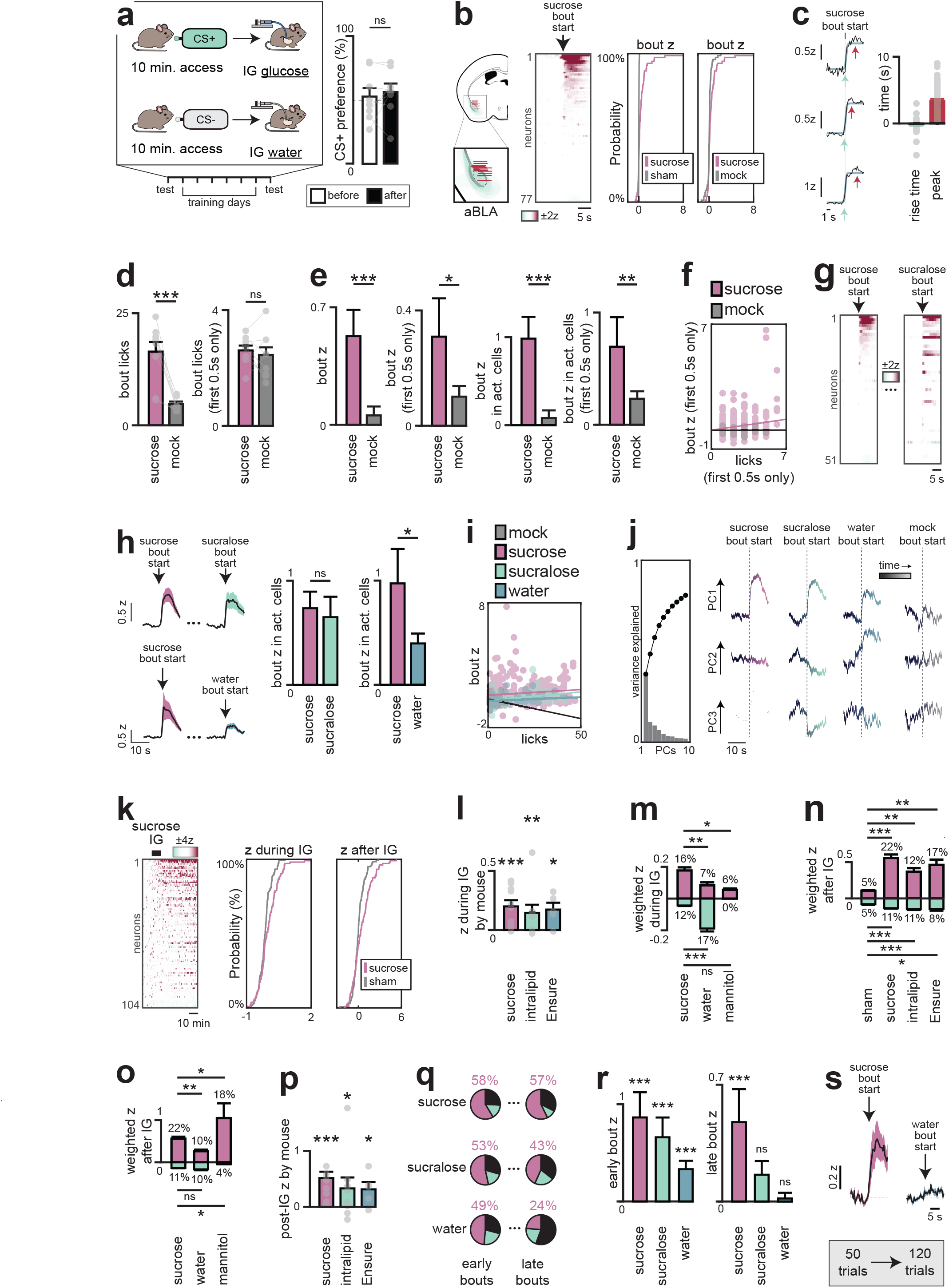
aBLA neurons track sweet taste and nutrient information to drive learning. a, Protocol for delayed flavour–nutrient conditioning. Mice received 10 min access to a flavoured saccharin solution followed by IG infusion of water or glucose. Mice did not learn a preference if glucose infusion occurred only after licking ceased. b, Fiber placement and viral spread for aBLA recordings. Averaged aBLA neuron dynamics during sucrose bouts (z-score) and distributions of activity compared to sham (p = 6.7 × 10-15, KS test) or mock lick bouts (p = 0.005, KS test). c, Peak (magenta) and 5% rise times (mint) with example cells (left) and summary plots (right). d, Lick count across trials and during the first 0.5 s of bouts. e, Activity of all cells (left) and sucrose-activated cells (right) across trials and during the first 0.5 s of bouts. f, Correlation of activity and lick count during first 0.5 s of sucrose vs mock bouts (sucrose: R² = 0.030, p = 0.0017; mock: R² = 0.00014, p = 0.95). g, Cross-registered neuron dynamics during sucrose (left) and sucralose (right) bouts. h, Mean traces (left) and activity (right) of sucrose-activated neurons during sucrose vs sucralose and sucrose vs water bouts. Minor differences in sucrose activity reflect cross-registration. i, Correlation of activity and lick count during mock, sucrose, sucralose, and water bouts (mock: R² = 0.062, p = 0.21; sucrose: R² = 0.018, p = 0.016; sucralose: R² = 0.17, p = 1 × 10-7; water: R² = 0.076, p = 0.0089). j, PCA of cross-registered neurons across sucrose, sucralose, and water sessions: scree plot (left) and first three PCs during each bout (right). k, Single-neuron dynamics (left) and distribution of activity (right) during and after sucrose IG infusion (both: p = 0.002, KS test). l, Population-weighted z-score activity during IG infusions, averaged by mouse. m, Nutrient specific activation during IG infusion of sucrose, but not water or mannitol. n, Activity after IG infusions. o, Unlike early activation, post-IG activity is not nutrient-selective. p, Population-weighted z-score activity after IG infusions, averaged by mouse. q, Distribution of neurons activated (red), inhibited (blue), or unchanged (black) during early (bouts 2–6) vs late (last five) sucrose bouts. Percent activated given in red. r, Late sucrose bouts (but not sucralose or water) activate these neurons. s, Trial-based drinking sessions extended to 120 trials in a new cohort, which showed nutrient-specific bout responses. ns P > 0.05; *P < 0.05; **P < 0.01; ***P < 0.001. Shaded areas in mean traces and error bars in bar plots show SEM. See Supplementary Table 3 for statistical tests, n, and p-values.

**Extended Data Fig. 9.**
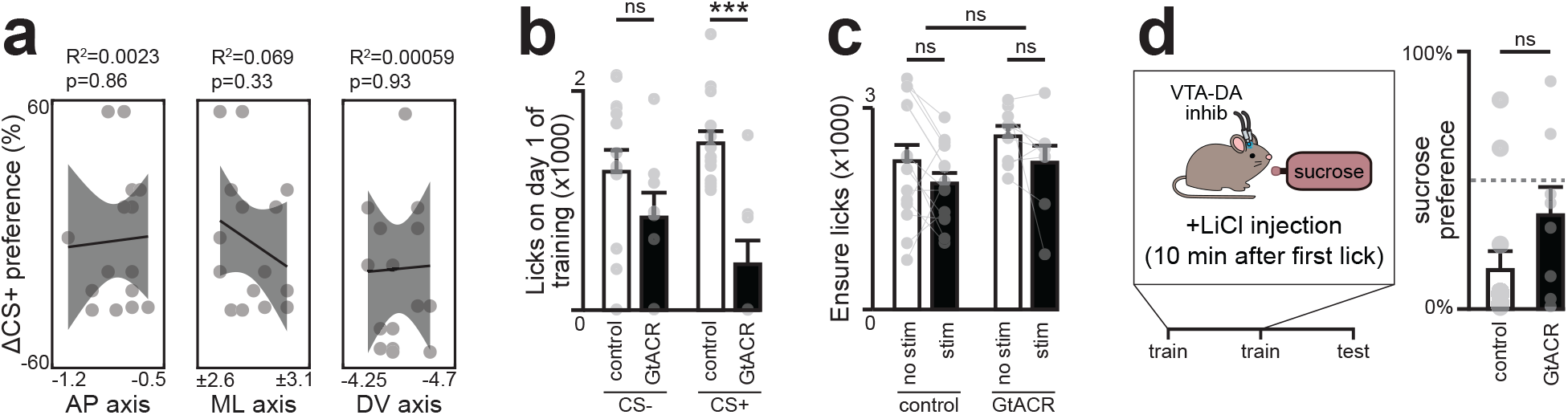
aBLA neurons are necessary for flavour-nutrient learning. a, Preference differences were not explained by fiber localization across the three axes. b, aBLA-D1R inhibition reduced CS+ intake on the first training day. c, Continuous inhibition did not alter consumption of the familiar nutritive solution Ensure. d, Conditioned taste aversion assay; inhibition during training did not block LiCl-induced aversion learning. ns P > 0.05; *P < 0.05; **P < 0.01; ***P < 0.001. Shaded areas in correlation plots and error bars in bar plots show SEM. See Supplementary Table 1 for statistical tests, n, and p-values.

**Extended Data Figure 10.**
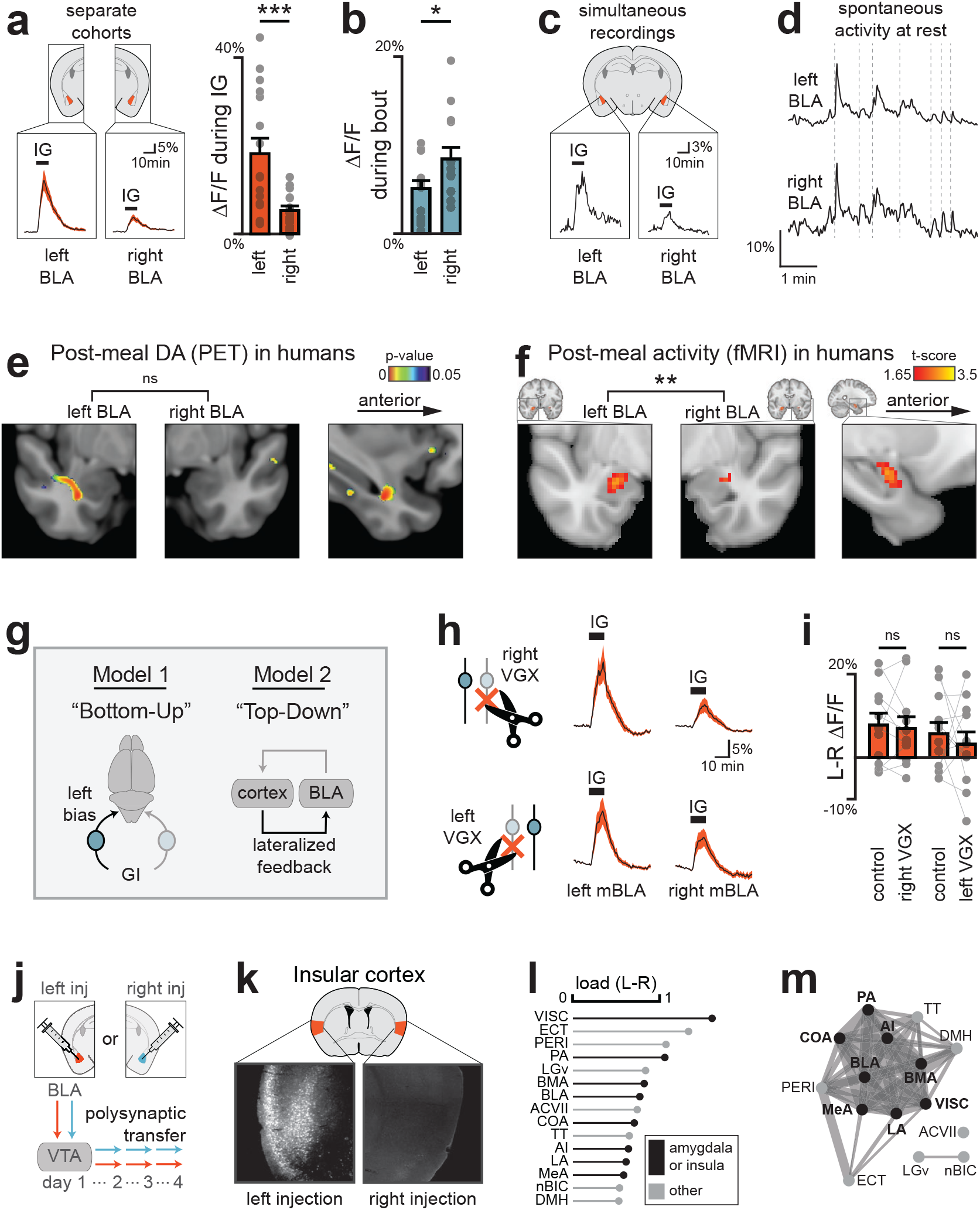
Internal nutrients preferentially engage the left BLA. a–b, Unilateral recordings in left or right BLA. a. Mean DA release during Ensure IG infusion is greater in left BLA (left, mean traces; right, summary plots). b, DA release during the first lick bout is greater in right BLA. c–d, Simultaneous recordings in left and right BLA. c. Example from one mouse showing DA activity in left and right BLA during Ensure IG infusion. d, Example from one mouse at baseline. e, In humans, PET (11C-raclopride) shows significant DA release after milkshake in left BLA but not right BLA (left-right difference not significant). f, fMRI shows significantly stronger activation after glucose consumption in the left relative to the right BLA. g, Left dominance may arise from (1) asymmetric vagal input, or (2) lateralized feedback circuitry. h, Bilateral mBLA dopamine recordings during Ensure IG after unilateral vagotomy (VGX). i, Difference in DA release between left and right mBLA before and after VGX. j, Polysynaptic retrograde tracing mapped inputs to left vs right VTA→BLA dopamine pathways. k, Greater labelling of insular cortex following polysynaptic tracing from left pathway. l, Top fifteen regions showing greater labeling after left vs right BLA injection (normalized to VTA labeling; n = 8 left injections, 6 right injections). m, Force-directed graph illustrating strong connectivity among regions in (l). Mice were sated for all infusion recordings (here and elsewhere), unless otherwise stated. Brains shown in neurological orientation (left = subject’s left). Full list of abbreviations in Ex. Fig. 12. ns P > 0.05; *P < 0.05; **P < 0.01; ***P < 0.001. Shaded areas in mean traces and error bars in bar plots show SEM. See Supplementary Table 2 and 4 for statistical tests, n, and p-values. See Supplementary Table 5 for raw data from pseudorabies experiment. See Supplementary Table 6 for connectivity matrix.

**Extended Data Figure 11.**
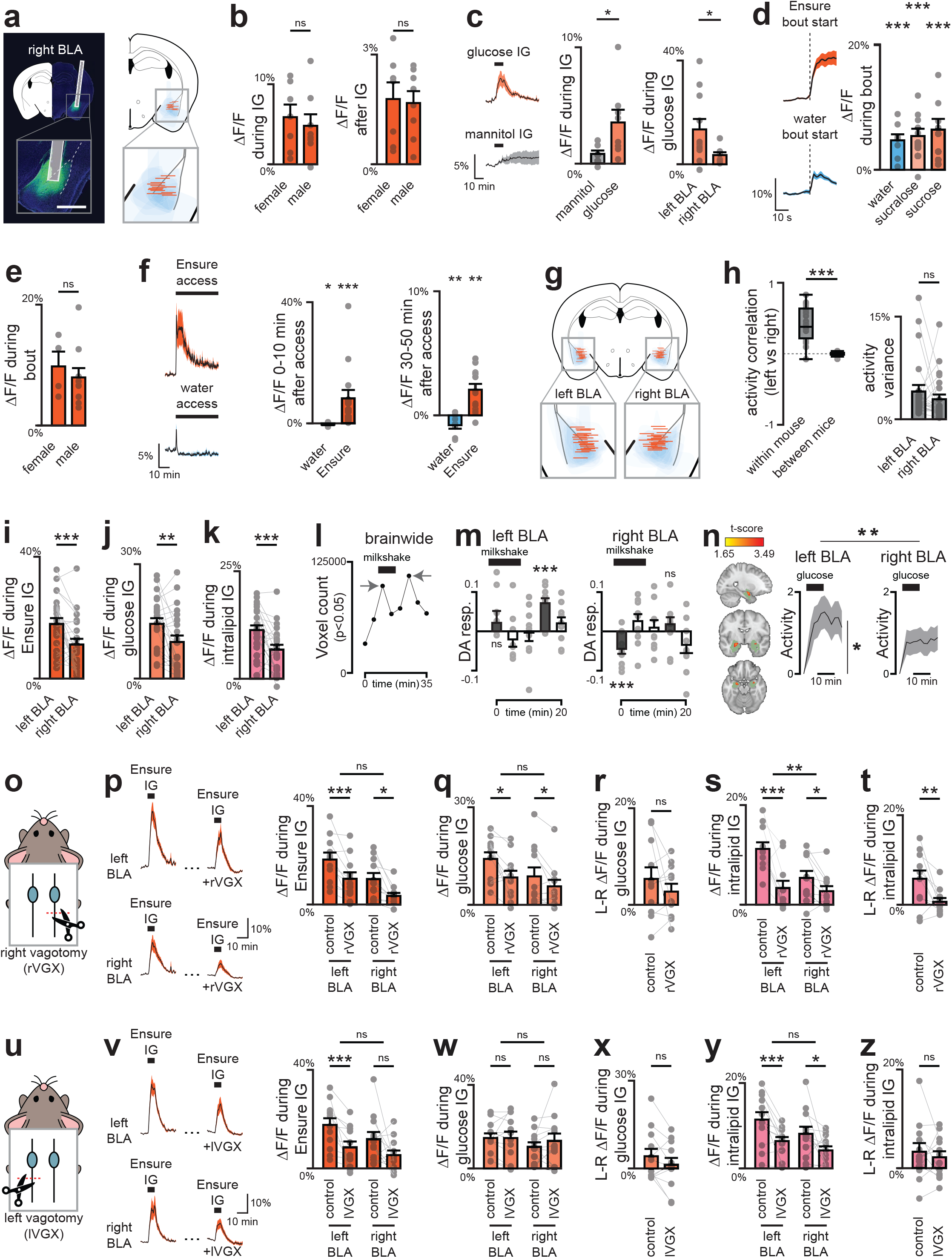
Nutrient-triggered dopamine release in the right BLA. a–f, Unilateral right BLA recordings. a, Fiber placement and viral spread for GRAB-DA recording in right BLA (scale bar, 1 mm). b, No sex differences in right BLA DA release during (left) or after (right) Ensure infusion. c, Mean traces (left) and summary plots showing glucose responses in right BLA are nutrient-specific (middle) but smaller than in left BLA (right). d, (Left) Mean trace showing DA release in right BLA during first bout of Ensure or water consumption. (Right) Summary plots of right BLA DA release during first lick bouts. e, No sex differences in right BLA DA release during first Ensure lick bout. f, Traces (left) and summary plots (right) of right BLA DA release during Ensure or water consumption. g–k, Bilateral BLA recordings. g, Fiber placement and viral spread for simultaneous GRAB-DA recording in left and right BLA. h, Spontaneous activity is coordinated (left) and comparable (right) across hemispheres. i, Ensure responses are larger in left BLA. j, Glucose responses are larger in left BLA. k, Intralipid responses are larger in left BLA. l–n, PET and fMRI recordings in humans. l, Number of brain-wide voxels (PET) showing increased dopamine release after milkshake consumption (normalized to water), with two peaks: an immediate oral signal (0–5 min after drinking onset) and a delayed post-ingestive signal (5–10 min after drinking offset). m, Time course of dopamine release in left and right BLA. Significance shown for oral and post-ingestive time bins (grey bars). n, Left and right BLA masks (left) and fMRI activity following glucose consumption (right). Activity was significant in left, but not right BLA, and direct comparison showed a significant difference between left and right BLA (F(1,703) = 7.83, *p* = .005). Green overlay refers to anatomically defined BLA masks by Amunts et al., 2020. o–z, Vagotomy experiments. o, Right vagotomy (rVGX). p, Ensure responses in both hemispheres partially depend on right vagus. q, Glucose responses in both hemispheres partially depend on right vagus. r, Right vagotomy does not affect lateralized glucose responses. s, Left BLA intralipid responses show greater sensitivity to right vagotomy. t, Right vagotomy abolishes lateralized intralipid responses. u, Left vagotomy (lVGX). v, Ensure responses in both hemispheres partially depend on left vagus. w, Left vagotomy does not affect glucose responses in either hemisphere. x, Left vagotomy does not affect lateralized glucose responses. y, Intralipid responses in both hemispheres partially depend on left vagus. z, Left vagotomy does not affect lateralized intralipid responses. ns P > 0.05; *P < 0.05; **P < 0.01; ***P < 0.001 (FWE-corrected for human data). Shaded areas in mean traces and error bars in bar plots show SEM. In box plots, top and bottom edges define 25th and 75th percentile; whiskers show maximum and minimum values. See Supplementary Table 2 and 4 for statistical tests, n, and p-values.

**Extended Data Figure 12.**
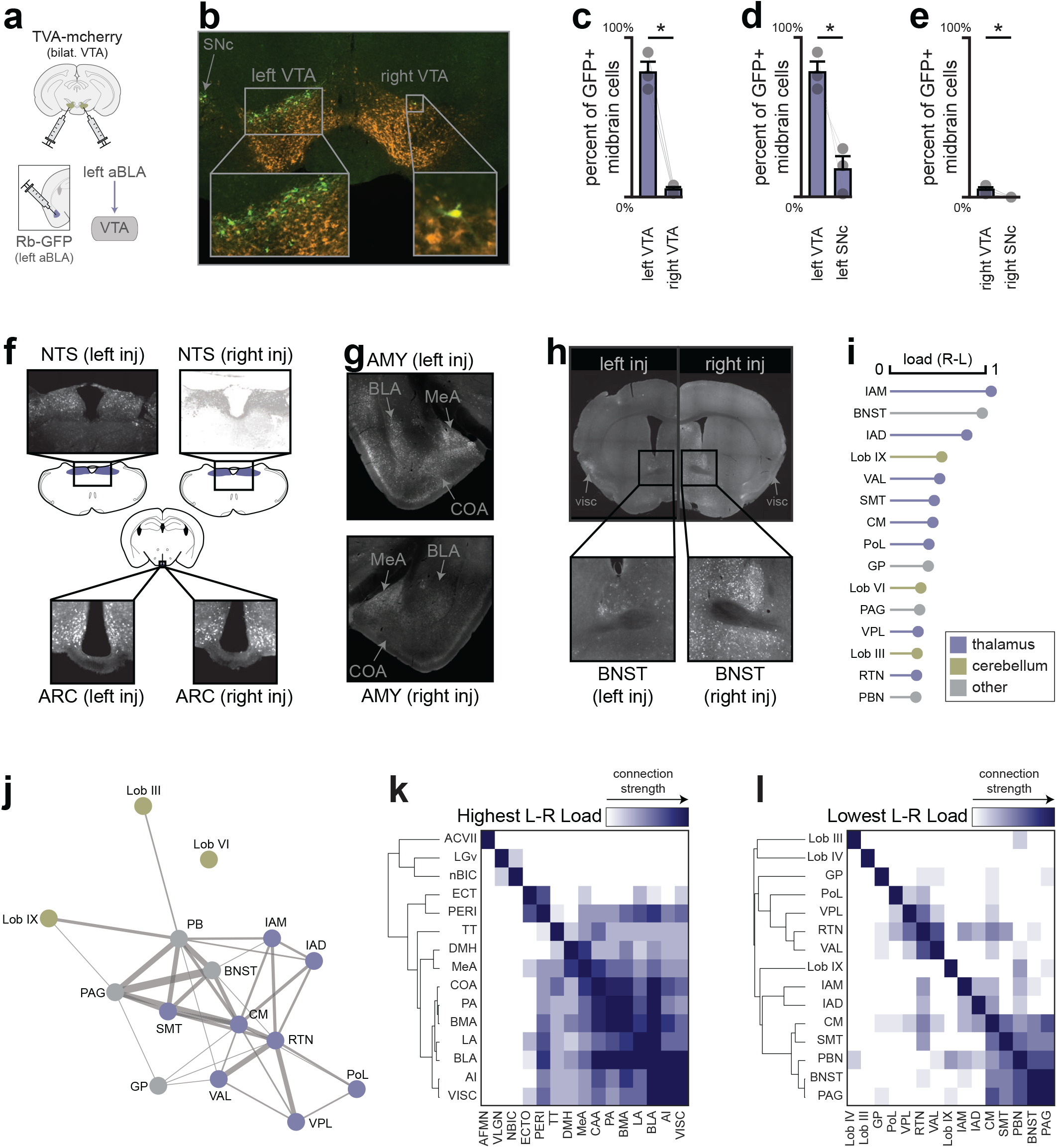
Networks underlying dopamine lateralization. a, Rabies tracing of DAT+ midbrain neurons projecting to left aBLA. Cre-dependent TVA was expressed in VTA/SNc, followed 3 weeks later by injection of CVS-N2c-dG-mGFP in left aBLA. b, Labelled midbrain cells 1 week after rabies injection, showing strong ipsilateral bias. Insets show a single contralateral VTA cell; some labeling in ipsilateral SNc. c–e, Quantification of labeled cells: 78 ± 7% ipsilateral VTA, 17 ± 8% ipsilateral SNc, 5 ± 2% contralateral VTA, and none in contralateral SNc. f, Cells labelled four days after PRV-Introvert-GFP injection into left BLA and right BLA of DAT-cre mice. No substantial differences observed in the nucleus of the solitary tract (NTS) or arcuate nucleus (ARC), where nutrient signals enter the brain. g, Greater labelling of several amygdalar regions after left-side injection. h, Greater labelling of BNST following right vs left injection (arrows indicate VISC showing greater labelling after left injection). i, Top fifteen regions that demonstrated strongest labelling after injection into right vs left BLA. j, Force-directed graph showing connectivity between areas in (e). k–l, Connectivity matrixes and hierarchical clustering of areas with greatest labelling after injection into left vs right BLA (f) or right vs left BLA (g). Abbreviations (used here and Ex. Fig. 10): ACVII, accessory facial motor nucleus; AI, agranular insula; ARC, arcuate nucleus; BLA, basolateral amygdala; BMA, basomedial amygdala; BNST, bed nuclei of the stria terminalis; CM, central medial nucleus of the thalamus; COA, cortical amygdala area; DECLIVE, lobule VI (declive); DMH, dorsomedial nucleus of the hypothalamus; ECT, ectorhinal area; GP, globus pallidus; IAM, interanteromedial nucleus of the thalamus; IAD, interanterodorsal nucleus of the thalamus; LA, lateral amygdala; LGv, ventral lateral geniculate; Lob III, lobule III; Lob IX, uvula; MeA, medial amygdala; nBIC, nucleus of the brachium of the inferior colliculus; NTS, nucleus of the solitary tract; PA, posterior amygdala; PAG, periaqueductal gray; PBN, parabrachial nucleus; PERI, perirhinal area; PoL, posterior limiting nucleus of the thalamus; RTN, reticular nucleus of the thalamus; SMT, submedial nucleus of the thalamus; TT, taenia tecta; VAL, ventral anterior-lateral complex of the thalamus; VISC, visceral area; VPL, ventral posterolateral nucleus of the thalamus. ns P > 0.05; *P < 0.05; **P < 0.01; ***P < 0.001. Error bars in bar plots show SEM. See Supplementary Table 5 for raw data from pseudorabies experiment. See Supplementary Table 6 for connectivity matrix.

