## Supplemental Text for "The neural basis of flavour-nutrient learning"

**Description of Extended Data Tables**

**Extended Data Table 1. Behavior statistics.** Summary information for all behavioral tests for each manipulation. Values are reported to 1 decimal place. Number of animals is reported as n. P-values are reported for all statistical tests, and are shown in bold if below standard significance level (α=0.05). In cases where a Benjamini-Hochberg correction was made for multiple comparisons, an adjusted p-value (p’) is shown (where p’=p*m/i, where i is rank and m is number of cohorts tested; see Methods). X’ corresponds to the normalized value of X. For self-stimulation, ‘presses means the number of inactive lever presses have been subtracted from the number of active lever presses. In flavor conditioning experiments where mice began the test with >90% preference for one flavor, statistics are also provided (in parentheses) with these mice excluded. Permutation (perm.) tests use an exact test (or 100,000 iterations if n>9) and are unpaired, unless otherwise mentioned (note: this means the lowest calculable p is 1E-5).

**Extended Data Table 2. Dopamine imaging statistics.** Summary information for all recordings of dopamine release in each brain region. Values are reported to 2 decimal places. Number of animals is reported as n. P-values are reported for all statistical tests, and are shown in bold if below standard significance level (α=0.05). X’ corresponds to the normalized value of X. For photometry traces, dF’/F’ means the dF/F values seen during sham infusion have been subtracted. For rise times, s’ means the time is relative to approximate entry of liquid into the stomach. For fall times, s’ means the time is relative to the end of the infusion. For peak times during licking, s’ means the time is relative to the start of the first licking bout. Baseline is defined as the first 10 minutes of the recording for GI and post-GI measurements, or 0–20 seconds before the start of the first licking bout for oral measurements. Permutation (perm.) tests use an exact test (or 100,000 iterations if n>9) and are unpaired, unless otherwise mentioned (note: this means the lowest calculable p is 1E-5).

**Extended Data Table 3. Single cell imaging statistics.** Summary information for all 1 and 2 photon miniscope recordings. Values are reported to 2 decimal places. Number (n) of animals and number of cells are reported separately. P-values are reported for all statistical tests, and are shown in bold if below standard significance level (α=0.05). X’ corresponds to the normalized value of X. For rise times, s’ means the time is relative to either the onset of the licking bout or to the approximate entry of liquid into the stomach. Population-weighted activity (pop-z) is calculated for a group of neurons (e.g. activated or inhibited neurons) by multiplying activity by the fraction of neurons belonging to that group. Linear-mixed models (LME) treat mouse differences as both a fixed random effect and covariate (see Methods). The relative contribution of mouse differences to total variance is reported by intra-class correlations (ICC). Note that for PCA, the number of PCs is equal to the number of cells. Permutation (perm.) tests use an exact test (or 100,000 iterations if n>9) and are unpaired, unless otherwise mentioned (note: this means the lowest calculable p for those tests is 1E-5).

**Extended Data table 4. fMRI imaging statistics (n=38).** Activity from left and right BLA clusters binned every 2 minutes (20 min total), as shown in Fig. 10f and Extended Fig. 11n. For PET statistics, see Thanarajah et al., 2019. Each of the 10 time bins after glucose intake was contrasted against the baseline (2 min before glucose intake), representing the change in brain activity after glucose intake relative to baseline. The contrast for the first effect was [ones(1,10)], and for the following contrasts [1], corresponding to the different time bins. Significance threshold was p < .05, FWE-corrected for multiple comparisons at the voxel level within the left or right BLA mask, respectively, with initial peak-level threshold set to p < .05 uncorrected.

**Extended Data table 5. Pseudorabies statistics.** Regional labeling (“load”) was normalized to VTA load to control for injection efficiency. From the 1,328 annotated CCFv3 regions, only gray-matter areas were retained (372 total), with cortical layers collapsed into single structures. Regions smaller than 100 × 100 × 100 pixels were excluded to remove small nuclei with high variance from few labeled cells, yielding 131 analyzed regions.

**Extended Data Table 6. Brain connectivity tables.** Connectivity matrices were generated manually, with each pair given a number indicating bidirectional projection strength from 0 (no known direct connection) to 5 (strongest projection known from this area). The connectivity strength between an area and itself was assigned a value of 5. For deciding what strength to assign a projection, large databases like the Allan Mouse Brain Connectivity Atlas^1^ were preferred, supplemented by additional sources comprising reviews and anatomical tracing studies^2–32^.
